# PhenoMapR: scalable mapping of sample phenotypes to single-cell, spatial, and bulk transcriptomics data

**DOI:** 10.64898/2026.09.08.749933

**Authors:** Brooks A. Benard, Chinmay K. Lalgudi, Armon Azizi, Andrew J. Gentles

**Affiliations:** Department of Pathology, Stanford University, Stanford, CA, 94035, USA; Department of Biochemistry, Stanford University, Stanford, CA, 94035, USA; Department of Internal Medicine, University of California San Diego, La Jolla, CA, 92093, USA; Department of Medicine, Stanford University, Stanford, CA, 94035, USA; Department of Biomedical Data Science, Stanford University, Stanford, CA, 94035, USA

## Abstract

Single-cell and spatial transcriptomic studies often lack sufficient sample size to compute robust statistical associations between a sample-level phenotype and cell types or spatial locations. In contrast, lower resolution methods such as bulk gene expression profiling have been applied at scale in large, annotated datasets, providing reliable signatures for phenotype associations. We introduce PhenoMapR, a semi-supervised method designed to integrate the phenotypic rigor of large-scale bulk expression studies with the cellular and spatial granularity of single-cell and spatial transcriptomics. PhenoMapR achieves this by deriving and mapping bulk gene expression signatures onto cells and spatial locations in a computationally efficient and scalable manner. The framework is broadly applicable across biological contexts, supporting the mapping of binary, continuous, and survival phenotypes derived from bulk expression studies across transcriptomic data modalities. This enables the identification of biologically-relevant cellular populations and spatial niches for experimental validation and therapeutic intervention.

## INTRODUCTION

Gene expression profiling from bulk tissue samples is one of the most common ways to efficiently measure and robustly stratify patients across cancers and other disease indications. Compared to other modalities such as mutations, methylation, protein expression etc., gene expression profiling remains one of the most predictive measurements for assigning prognosis across cancers^1–3^. Prior work has helped to refine the interpretability of bulk gene expression data. For example, there are now multiple methods to reliably deconvolve bulk gene expression into cell type proportions^4,5^, refined high-resolution cell-type signatures^6^, and co-existing cellular ecosystems and states^7,8^. These approaches have been applied at scale and have helped to define our understanding of the impact of single genes^9–11^, gene signatures^3^, cell type proportion^10,12^, and cellular ecosystems^7^ on cancer prognosis and response to therapy.

While bulk sequencing approaches have proven powerful for large-scale associations they remain limited by the biological complexity collapsed in each sample, constraining the mechanistic understanding and hypothesis generation possible from these methods. The development and application of single-cell and spatial transcriptomic (ST) technologies across biological fields has dramatically improved the resolution of expression-based measurements^13^. These methods have enabled unprecedented mapping of gene expression signatures to individual cells and spatial locations in tissues. Despite these advances, single-cell and ST approaches remain far more expensive than bulk gene expression profiling. Due to this constraint, many single-cell and ST studies lack the sample size required to identify robust statistical associations between sample phenotypes (e.g. mutation status, outcomes, etc.) and cell types or locations^14–18^. To bridge this gap, approaches have emerged to identify cellular and spatial origins of phenotypic signatures to improve resolution and identify therapeutic targets^19–21^.

Several methods have been developed to identify phenotypically-relevant cells and locations from single-cell and spatial data^22–32^. These approaches have helped to transfer the wealth of information derived from bulk expression datasets to single-cell and spatial resolution. However, these methods generally do not scale well and often run into time and compute limitations, restricting their application to increasingly large datasets and atlases^33^. Additionally, all existing methods require as input a phenotype-labeled bulk expression dataset, placing the onus on users to identify, obtain, and preprocess an appropriate reference. This process is time-consuming, introduces analytical variability, and can present a significant barrier to use.

To address these limitations, we developed PhenoMapR, a scalable and flexible framework for identifying cells, locations, and samples from high-resolution gene expression data that are associated with sample phenotypes in bulk transcriptomics datasets. PhenoMapR comes pre-loaded with 138 pan-cancer adult, pediatric, and immunotherapy outcome signatures for fast rank-ordering of samples, cells, and locations most associated with favorable and adverse prognosis. Additionally, PhenoMapR can preprocess bulk expression reference datasets for custom phenotype signature generation, facilitating its application in targeted datasets and non-cancer settings. Here, we demonstrate PhenoMapR’s utility by performing pan-cancer bulk expression outcome stratification with targeted analyses in pancreatic adenocarcinoma (PAAD) single-cell and spatial data, as well as single-cell analyses in non-cancer contexts, including COVID-19, Alzheimer’s disease (AD), and facioscapulohumeral muscular dystrophy (FSHD). Across all expression modalities and disease indications, we recapitulate known biology while also identifying novel associations for future validation. Additionally, we demonstrate order-of-magnitude improvements in both runtime and memory usage requirements compared to other tools, while retaining sensitivity and specificity of results. The PhenoMapR package and interactive Shiny app are available at https://github.com/brooksbenard/PhenoMapR.

## RESULTS

### Development and benchmarking of PhenoMapR

PhenoMapR is a semi-supervised approach for identifying cells, locations, and samples that express transcriptional signatures associated with a sample phenotype (**Fig. 1a**). Given a bulk expression dataset containing sample-level annotations, PhenoMapR derives gene-level phenotype association z-scores using either logistic regression (binary labels), correlation (continuous labels), or Cox proportional hazards regression (time-to-event labels). Then, for each bulk, single-cell, or spatial transcriptomics input, PhenoMapR computes the weighted-sum between the reference phenotype signature and the normalized expression matrix (**Fig. 1b**). The resulting scores thus take into account both the relative expression levels of individual genes as well as their phenotypic weight. Additionally, PhenoMapR contains pre-computed prognostic meta-Z-score signatures for all cancers in The Cancer Genome Atlas (TCGA)^34^ and PRECOG^10^ (adult, pediatric, and immunotherapy cohorts), facilitating immediate scoring of an input cancer dataset without the need for manually deriving a bulk phenotype signature (**Fig. 1c**). This built-in reference database reflects the largest curation, annotation, pre-processing, and survival outcomes signature generation across cancers to-date: 335 datasets, >46,000 patients, and 55 cancer subtypes^10^.

**Fig. 1:**
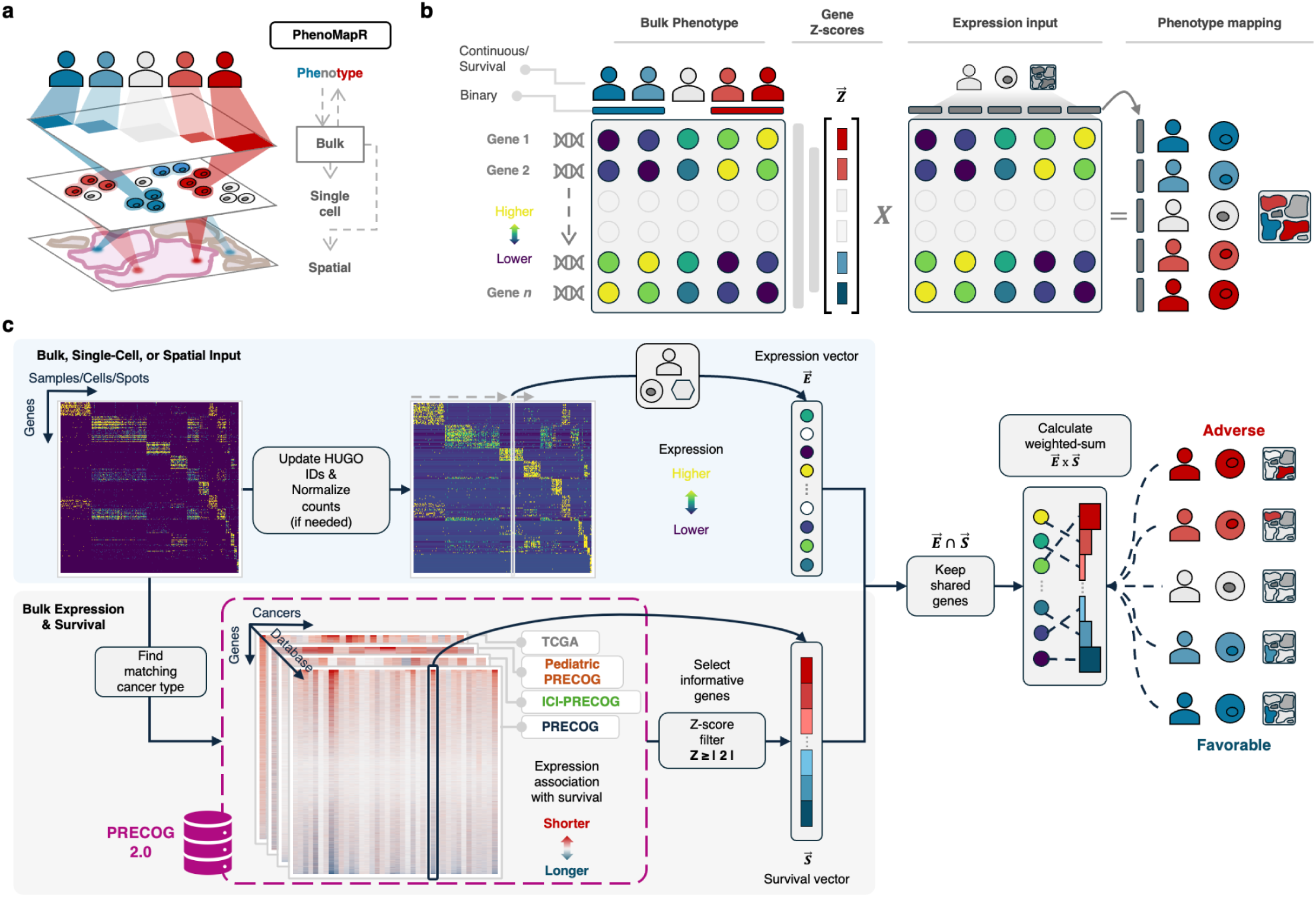
Overview of PhenoMapR. **(a)** Schematic representation of mapping sample phenotypes across transcriptional modalities of different scales. **(b)** High-level conceptualization of PhenoMapR; bulk expression data with sample phenotype annotations can be used to derive a gene z-score signature which can then be used to identify phenotype-associated samples/cells/locations. **(c)** Diagram of scoring against built-in, pre-computed phenotype references from PRECOG and TCGA.

To benchmark PhenoMapR, we implemented SigBridgeR^35^, a framework and comprehensive toolkit that unifies phenotype-association algorithms into a consistent workflow for accurate comparison **(Supplementary Fig. 1a)**. We performed runtime and memory scaling analyses across eleven other methods and observed that PhenoMapR showed substantially better performance across both metrics **(Supplementary Fig. 1b)**. Additionally, we compared cell-level results across all methods with PhenoMapR results using either enrichment or correlation analysis **(Supplementary Fig. 1c-e)**. Across almost all methods, phenotype-associated cells showed strong enrichment at the tails of PhenoMapR score distributions. Comprehensive cross-method comparisons were also performed (**Supplemental Fig. 1f,g**). Overall, PhenoMapR emerged as orders-of-magnitude more efficient than other methods, scaled efficiently with dataset size, and showed strong concordance with the vast majority of other established algorithms.

### Application of PhenoMapR in stratifying bulk expression datasets

We first tested if PhenoMapR could recover gene-phenotype associations derived from bulk data in a separate bulk dataset of the same disease type. Bulk expression signatures associated with outcomes across cancers have been thoroughly described^3,10,36,37^. To demonstrate the feasibility of using PhenoMapR for assigning phenotype signatures (i.e. survival) at the sample level, we processed cancers from TCGA using built-in outcomes signatures from PRECOG (**Fig. 2a**). This was only performed for the 25 cancer types shared between PRECOG and TCGA (**Supplementary Fig 2a**). All bulk TCGA cancer samples were scored using PRECOG meta-z signatures, stratified by median sample score, and tested for significant differences in outcomes. In total, 22 of 25 cancers showed the expected directionality of hazard ratios, with 15 of 25 showing statistically significant differences in overall survival based on a median PhenoMapR score (**Fig. 2b**). Two cancer types (UCS and READ) showed an inverse trend in outcomes associations, perhaps related to small cohort and gene signature sizes. For the other non-significant cancers, the lack of significance could be explained by low TCGA vs. PRECOG meta-z correlation (e.g. DLBC, CESC, LUSC) and/or small sample size (e.g. DLBC). As a positive control, TCGA cancers scored against built-in TCGA meta-z signatures showed significant stratification in 27 of 33 cancer types (**Supplementary Fig. 2b**). These results demonstrate that PhenoMapR can be used to score and stratify bulk gene expression datasets in a pan-cancer approach.

**Fig. 2:**
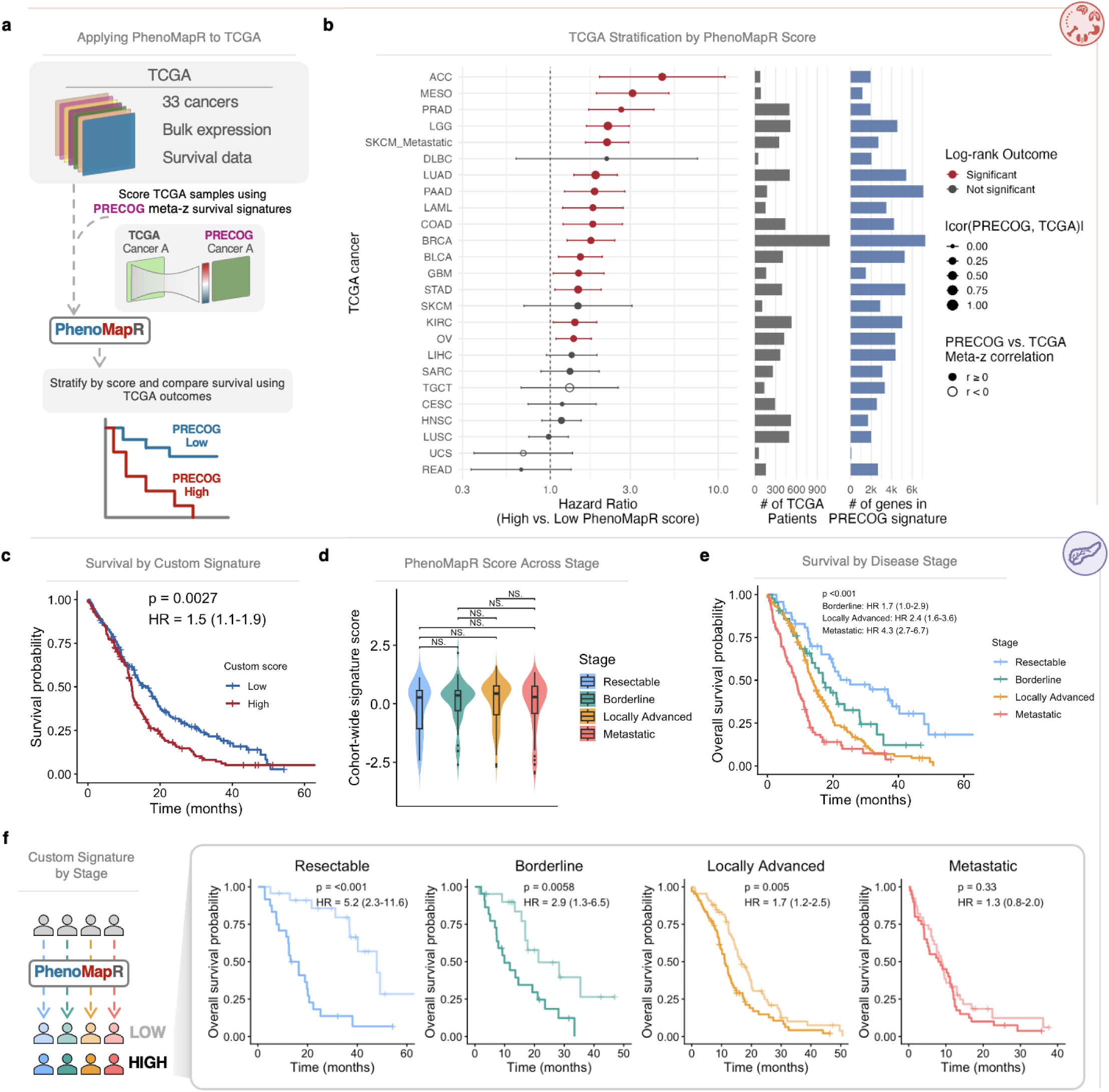
Application of PhenoMapR in bulk expression datasets. **(a)** Schematic of using PhenoMapR to score and stratify TCGA cancers for outcomes associations. **(b)** Hazard ratios for each TCGA cancer type when stratified by median PhenoMapR score (left). Number of patients per TCGA cancer type with both expression and outcomes data available (middle). Number of genes per cancer type with significant PRECOG survival associations (|Z ≥ 2|) used to score TCGA patients (right). **(c)** Survival for n=317 pancreatic adenocarcinoma (PAAD) patients based on a median PhenoMapR score split using custom scoring. **(d)** PhenoMapR score distribution across disease stages. Significance between mean distributions was calculated using a two-sided Wilcoxon rank-sum test. **(e)** Outcomes for the different disease stages in GSE253260. **(f)** Survival curves within each disease stage using a median split of stage-specific PhenoMapR-derived signature scores. P-values for survival were computed using a two-sided log-rank test. Hazard ratios were calculated using univariate Cox proportional hazards regression.

### Deriving a custom reference signature with PhenoMapR

PhenoMapR provides functionality to derive custom phenotype signatures based on input bulk expression and sample phenotype labels (binary, continuous, or survival outcomes). We demonstrate custom signature generation and application using a bulk expression dataset of 317 clinically-annotated PAAD patients across the disease stage continuum^38^. Using all samples for signature generation, the resulting PhenoMapR score was able to significantly stratify outcomes across the cohort but not within stages (**Fig. 2c and Supplementary Fig. 2c**). This was interesting because the PhenoMapR score was not significantly different across disease stages (**Fig. 2d**) yet the different disease stages had quite distinct outcomes (**Fig. 2e**), suggesting that PhenoMapR identified potentially stage-agnostic prognostic signals and might be able to refine the risk within each stage. Indeed, by generating stage-specific custom signatures, PhenoMapR was able to stratify outcomes for all stages except metastatic, with decreasing hazard ratios, C-indexes, and multivariate significance as disease stage advances (**Fig. 2f and Supplementary Fig. 2d,e**). These results show how custom signature generation with PhenoMapR can provide clinically-relevant prognostic insights in PAAD, potentially nominating a subset of traditionally lower-risk patients for more aggressive intervention.

### PhenoMapR identifies cell types associated with COVID-19 disease severity

The peripheral blood immune system correlates to clinical disease severity in COVID-19 have been extensively studied^39^, thus making it an ideal setting to test PhenoMapR. We performed positive control analyses in several large COVID-19 datasets to show that PhenoMapR correctly identifies phenotype-related cellular populations using both binary (healthy vs. disease) and continuous (disease severity) approaches. Across three large single-cell studies^40–42^ (>850,000 cells across 180 patients), we generated sample-level pseudobulk expression profiles using half of each cell type per sample. These pseudobulks were used to generate study-specific COVID-19 severity signatures using binary and continuous PhenoMapR approaches; held-out single cells were scored and cell-type associations were compared. Across all datasets, PhenoMapR successfully identified the reported monocyte (e.g. proliferating/CD14^+^) and neutrophil (e.g. immature/developing) populations as being associated with more severe disease, while natural killer cell subpopulations were correctly associated with more favorable disease (**Supplementary Fig. 3a-f**). This illustrates that PhenoMapR is able to recover known biology of COVID-19 across independent cohorts and confirms strong correlation between binary and continuous phenotype assignment approaches.

### PhenoMapR nominates clinically-actionable subpopulations in single-cell PAAD data

To demonstrate PhenoMapR’s use in the context of cancer, we leveraged a large dataset of 57,443 cells from 24 primary PAAD tumors and 11 normal pancreases^43^ (**Fig. 3a; Supplementary Fig. 4a**). We used PhenoMapR’s built-in TCGA and PRECOG PAAD overall survival outcome meta-z signatures to score individual cells, observing robust correlations between reference signatures (**Supplementary Fig. 4b**).

**Fig. 3:**
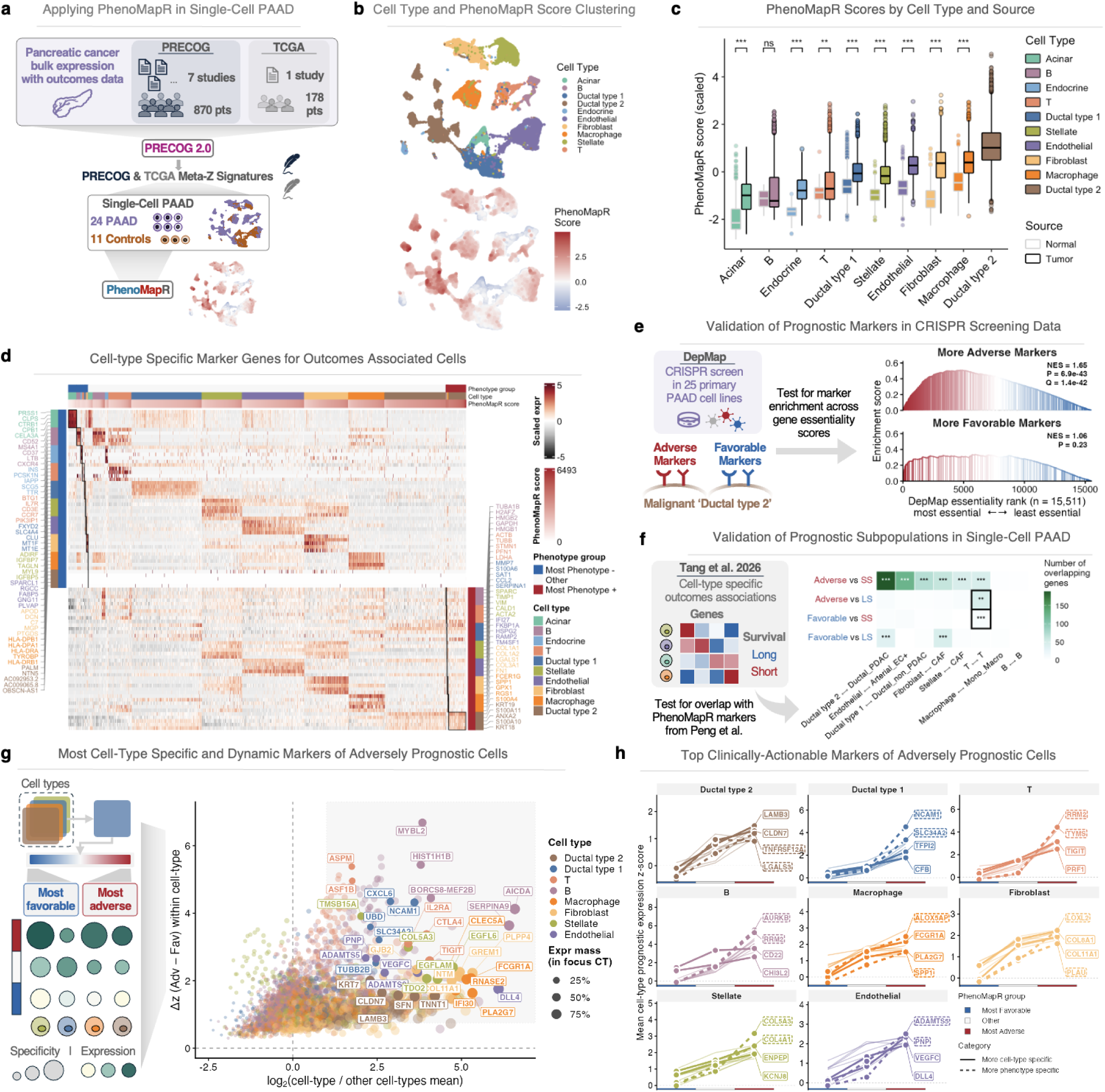
PhenoMapR nominates cell-type specific targets in single-cell PAAD. **(a)** Applying PhenoMapR to score single-cell PAAD data using PAAD outcomes meta-z signatures from PRECOG 2.0. **(b)** UMAP representation of cell types (top) and scaled PRECOG-derived PhenoMapR scores (bottom) for cells in the Peng et al.^43^ single-cell dataset. **(c)** PhenoMapR score distribution across cell types and sample source (tumor/normal) ordered by increasing mean cell type score. **(d)** Heatmap summary of the top cell-type specific marker genes for the most adverse and favorable cell populations in the dataset (5th percentile). “Phenotype +” labels reflect cells associated with worse outcomes and “Phenotype -” labels indicate cells associated with more favorable outcomes. Black boxes indicate marker genes for the cell-type at a phenotype extreme. Some cell types don’t enter the 5% filter. **(e)** Gene set enrichment analysis of marker genes associated with the most adverse/favorable malignant ‘ductal type 2’ cells across PAAD cell lines in the DepMap database. **(f)** Overlap of cell-type specific prognostic genes from Tang et al. with prognostic marker genes derived by applying PhenoMapR in the Peng et al. dataset. Black boxes around heatmap tiles signify overlap of genes in opposite phenotype populations. LS = Longer overall survival; SS = Shorter overall survival. **(g)** Characterization of the most adversely prognostic cell-type markers within tumor samples. Grey box represents the heuristic thresholds for markers with the more cell-type specific (x-axis) and prognostic delta (y-axis) specificity. **(h)** Mean expression across PhenoMapR groups and cell types for top clinically actionable adverse markers from DGIdb. P-values by two-sided Wilcoxon rank-sum test. Stars use Benjamini–Hochberg FDR on two-sided p(|Z|): *** q≤0.001, ** q≤0.01, * q≤0.05

Malignant ‘ductal type 2’ cells showed the strongest association with an adverse phenotype while acinar and B cells were enriched for favorable scores (**Fig. 3b,c; Supplementary Fig. 4b,c**), consistent with previous reports^43,27^. As expected, across all cell types, cells from normal pancreases scored as significantly more favorable (**Fig. 3c**). Importantly, scoring across cell types was resilient to the effects of cell cycle, donor variation, reference signature permutation/source, and sequencing depth (**Supplementary Fig. 4d-k**).

PhenoMapR provides automated marker gene identification for cells at the tails of a dataset score distribution (default 5th percentiles). We applied PhenoMapR’s cell-type specific marker gene functionality to identify markers for cells most associated with PAAD survival, revealing clear trends in cell-type marker gene expression across the range of PhenoMapR scores (**Fig. 3d**). For example, favorable acinar cells express genes such as *PRSS1* and *CTRB1* (fully differentiated)^44^, favorable T cells express *CCR7* and *IL7R* (naive/central memory)^45^, while favorable fibroblasts uniquely express *C7* and *APOD* (complement-rich/adipose-stroma-like)^46^. In contrast, adverse T cells expressed genes like *TUBB* and *STMN1* (proliferating/exhausted) while adverse fibroblasts expressed genes like *LGALS1* and *FN1* (activated myofibroblastic CAF)^47,48^. Notably, activated T regulatory cells (*IL2RA*⁺/*FOXP3*⁺) co-expressing at least one canonical effector marker (*TIGIT*/*ICOS*/*CTLA4*/*CD39*) were almost exclusively observed in the most adverse T cell population (**Supplementary Fig. 4l**).

We validated these results using CRISPR screening data from 25 primary PAAD cell lines in DepMap^49^; in the malignant ‘Ductal type 2’ population, markers of the most adverse cells were significantly enriched for genes which had stronger dependency scores, while “more favorable” markers in this malignant population were not significantly associated with gene essentiality, as might be expected (**Fig. 3e**). Additionally, cell-type prognostic markers from PhenoMapR significantly overlapped cell-type specific survival-associated genes from an independent PAAD single-cell atlas (**Fig. 3f**). Of therapeutic significance, PhenoMapR identified many non-canonical targets in the tumor microenvironment (TME; **Fig. 3g and Supplementary Fig. 4m**). For example, *NCAM1* and *SLC34A2* (druggable with agents like Lorvotuzumab and TUB-040, respectively) were uniquely overexpressed in non-malignant adverse ductal cells compared to other cell types in the TME (**Fig. 3g,h and Supplementary Fig. 4m**). Overall, PhenoMapR identified cellular populations with known roles in disease pathology and nominated novel cell-type targets in the TME for therapeutic validation.

### PhenoMapR identifies spatial correlates of prognostic signal in PAAD

PhenoMapR can be used to assign phenotype scores to spots/cells in spatial transcriptomics (ST) data. We utilized a 10X Visium ST PAAD sample from HTAN to investigate if phenotype signals have spatial architectures (**Fig. 4a**). When applied at the spot-level, PhenoMapR identified distinct spatial clusters of adverse and favorable prognostic signals that were strongly co-localized (**Fig. 4b**). We next used CytoSPACE^21^ to map paired single-cell data onto the ST spots for increased resolution. Running PhenoMapR on the mapped

**Fig. 4:**
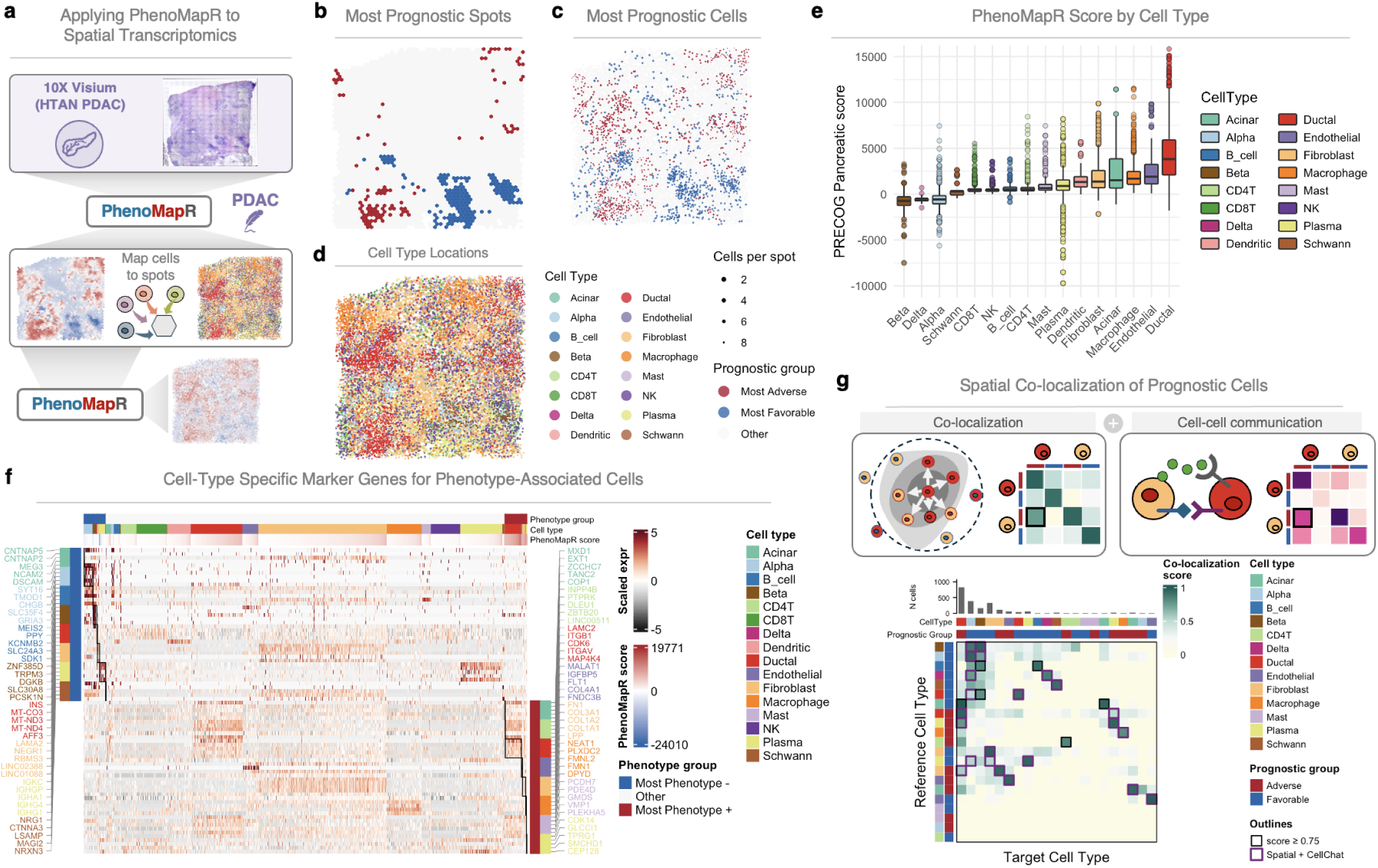
PhenoMapR identifies phenotypic signal in spatial transcriptomics data. **(a)** Schematic for processing spot-based expression data with PhenoMapR and then assigning single-cell level annotations and scores to spots. **(b)** Spot-based localization of the top and bottom 5th percentile of PhenoMapR scores. **(c)** Post-CytoSPACE PhenoMapR score of single-cell data. **(d)** Cell-type localization of CytoSPACE mapped cells. **(e)** PhenoMapR score distribution across cell types. **(f)** Cell-type specific marker genes between the most adverse and favorable cells based on PhenoMapR score. **(g)** Co-localization and cell-cell communication analyses grouped by cell type and prognostic group. Prognostic cell types showing both high co-localization scores and CellChat2 communication potential are indicated with a purple outline. single-cell data identified many more spatial locations with prognostic signal enrichment, while also confirming the broad co-localization of prognostic signal (Fig. 4c). Overlaying cell-type annotations revealed that ductal cells were the predominant source of adversely prognostic signal, while beta islet (and many plasma) cells contained the most favorably prognostic signal (**Fig. 4d,e**). We used PhenoMapR to identify marker genes for the most adverse phenotype-associated cells: ECM-high CAFs/stellate cells (*COL1A1*/*COL1A2*/*FN1*, *SPARC*/*ACTA2*/*VIM*), *SPP1*+ tumor associated macrophages, and malignant ductal type 2 programs (*KRT19*/*KRT18*/*ANXA2*) which aligns with established PDAC biology (Fig. 4f). Additionally, less canonical hits such as macrophage *RGS1*/*GPX1*, normal ductal *SAT1*, and endothelial *FKBP1A*/*RAMP2* emerged (Fig. 4f). Finally, we performed neighborhood/local co-occurrence enrichment^50^ and cell-cell communication analyses^51^ and observed distinct patterns of co-localization and communication between prognostic cells (**Fig. 4g, Supplementary Fig. 5a,b**). Specifically, prognostic cells tended to co-localize with other prognostic cells of the same type, however co-localization between differing cell types was also observed (**Fig. 4g, Supplementary Fig. 5a,b**). Of note, cell-cell communication analysis revealed autocrine laminin signaling between adverse ductal cells as well as collagen signaling between adverse fibroblasts and adverse ductal cells (**Supplementary Fig. 5b**). These results show that PhenoMapR can be used to identify phenotypically-associated cells in a spatially-relevant context.

### PhenoMapR identifies disease-relevant cells in Alzheimer’s Disease

To demonstrate PhenoMapR in a non-cancer setting, we leveraged bulk and single-cell datasets to identify cells associated with Alzheimer’s disease (AD) development. We used PhenoMapR to derive an AD phenotype score using bulk expression data from seven early AD and seven healthy controls^52^ and then used this signature to score single-cell data from six AD and six control samples^53^ (**Fig. 5a**). Across all cell types, AD samples scored worse than healthy control samples, with microglia, endothelial, and oligodendrocyte cell types enriched with the AD signature, while neurons contained more healthy control signal (while taking donor variation into effect, **Fig. 5b and Supplementary Fig. 6a**); this agrees with previous reports^23^. Microglia most associated with AD were marked by genes such as *DOCK4* and *PLXDC2*, both reported to be associated with AD progression and indicative of a phagocytically-impaired phenotype^54,55^. Healthy neurons had elevated expression of genes such as *ERBB4* and *NRXN3* and reduced expression or loss-of-function in these genes has been linked with the development of AD^56,57^.

**Fig. 5:**
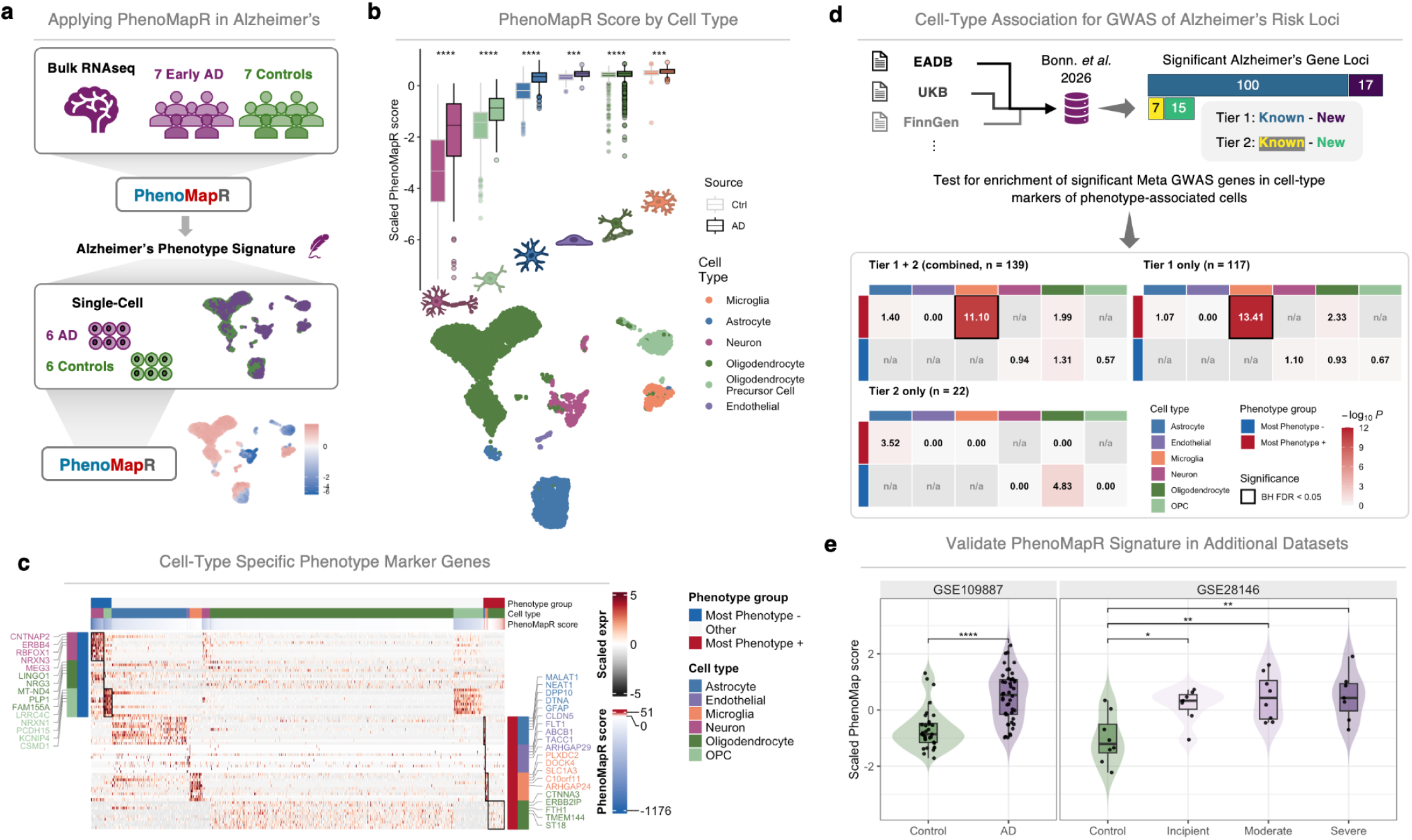
Application of PhenoMapR to Alzheimer’s disease. **(a)** Schematic of generating an Alzheimer’s phenotype signature with PhenoMapR and the signature’s application to a single-cell dataset using PhenoMapR. **(b)** Distribution of PhenoMapR scores across cell types and sample sources (top) and UMAP representation of cell types (bottom). **(c)** Heatmap of cell-type specific marker genes for cells in the 5th percentile of PhenoMapR scores. **(d)** Representation of meta GWAS results for AD gene loci (top) and cell-type enrichment of GWAS genes across phenotype groups (bottom). Heatmap values represent the odds ratio of GWAS genes being within the significant PhenoMapR marker genes for each cell type. Heatmap cells are n/a if a cell type was not present in the phenotype tail threshold. **(e)** Violin plots of scaled PhenoMapR scores across two different bulk gene expression studies using the PhenoMapR-derived AD signature from **a**. Each dot represents one sample. P-values by two-sided Wilcoxon rank-sum test; **** q≤0.0001, ** q≤0.01, * q≤0.05.

As an orthogonal validation of our findings, we leveraged results from a comprehensive meta-analysis of genome-wide association studies (GWAS) in AD^58^ to test for enrichment of AD-related GWAS genes in cell-type markers of phenotype-related cells (**Supplementary Fig. 6b,c**). Indeed, AD-associated microglial cells were significantly enriched for AD risk genes compared to other phenotype-related cells (**Fig. 5d**), similar to what was reported in the meta GWAS study. Of note, the “tier 2” loci genes in the meta GWAS study required “further external validation” due to not meeting significance after conditional analyses^58^. Interestingly, only the “tier 1” loci genes were significant in our analysis (**Fig. 5d**), suggesting that PhenoMapR identified high-quality phenotype-related cells in AD. Additionally, we used PhenoMapR to validate the bulk AD signature in two independent cohorts^59,60^ and found that AD phenotype scores stratified AD from controls and across AD staging (**Fig. 5e**). Together, these results illustrate PhenoMapR’s ability to uncover and clarify cellular determinants contributing to AD pathophysiology.

### PhenoMapR nominates cells-of-interest in facioscapulohumeral muscular dystrophy

We next applied PhenoMapR to identify phenotype-associated single cells from facioscapulohumeral muscular dystrophy (FSHD). We used bulk expression data from 27 FSHD samples and eight controls^61^ to derive a FSHD PhenoMapR signature. This signature was then used to identify FSHD associated cells from a single-cell dataset of four FSHD samples and two controls^62^ (**Fig. 6a**). As reported in other studies^63,64^, the canonical FSHD driver gene *DUX4* and its target genes were not expressed/showed no variance across samples and were not incorporated into the PhenoMapR signature for scoring FSHD (**Supplemental Fig. 7a,b**). Regardless, after assigning cell types, cells from FSHD samples consistently showed higher FSHD PhenoMapR scores (**Fig. 6b; Supplemental Fig. 7c,d**). Proliferating vascular smooth muscle cells (VSMCs) on average had the strongest FSHD signature, while senescent VSMCs were associated with a healthier phenotype (**Fig. 6b**). Cell-type specific analyses showed that elevated expression of genes like *CALM2* in proliferating VSMCs, *PTMA* in synthetic VSMCs, and *TAGLN* in vascular fibroblasts were associated with FSHD (**Fig. 6c**); these specific gene/cell-type associations exemplify potentially novel (*CALM2, PTMA*) and previously reported (*TAGLN*)^65^ associations with the disease.

**Fig. 6:**
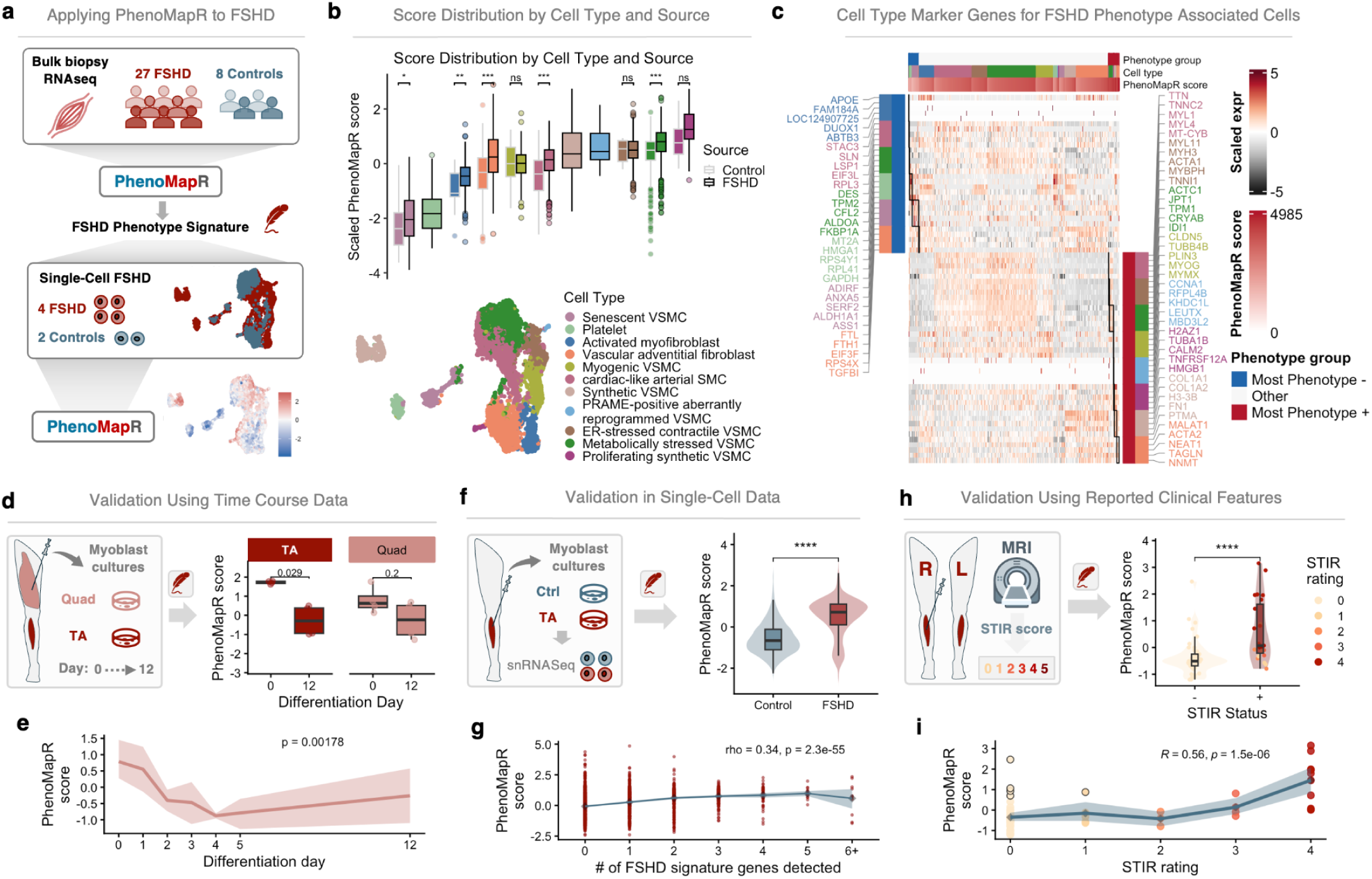
Identifying phenotype-associated cells in FSHD. **(a)** Generation of a FSHD phenotype signature from bulk expression data and application in a single cell dataset. **(b)** PhenoMapR score distribution by cell type and sample source. P-values by two-sided Wilcoxon rank-sum test; *** q≤0.001, ** q≤0.01, * q≤0.05, ns = Not Significant. **(c)** Heatmap of cell-type specific marker genes for cells in the 5th percentile of PhenoMapR scores. **(d)** Schematic (left) and PhenoMapR scores (right) of bulk FSHD samples from quadriceps (Quad) and tibialis anterior (TA) muscle biopsies cultured over a 12-day time course. P-values by two-sided Wilcoxon rank-sum test. **(e)** PhenoMapR score distribution for quadriceps samples across each timepoint of the 12-day time course. P-value calculated using a Jonckheere test. **(f)** Schematic (left) and PhenoMapR scores for control and FSHD TA single-cell myoblast cultures. **(g)** PhenoMapR scores (per single-cell) based on the number of FSHD signature genes detected per cell. Rho and p-values were calculated using a Spearman correlation. Diamonds represent the mean score per category. Ribbon shows the bootstrapped 95% CI per category. **(h)** Schematic (left) and PhenoMapR scores (right) of bulk bilateral FSHD TA biopsies stratified by STIR status. **(i)** PhenoMapR score versus STIR rating. Points outlined in black represent samples with discordant STIR rating and PhenoMapR scores. Rho and p-values were calculated using a Spearman correlation. Diamonds represent the mean score per category. Ribbon shows the bootstrapped 95% CI per category.

FSHD affects the upper body and tibialis anterior (TA) muscles earlier and more aggressively than other muscles such as quadriceps (quad)^66^. Using bulk RNAseq of paired FSHD samples from quad and TA muscle biopsies cultured ex-vivo over a 12-day time course^67^, PhenoMapR detected a significant decrease in disease signature between the two TA biopsies (**Fig. 6d**). Although not significant between time point extremes (i.e. Day 0 and 12), the quad PhenoMapR score did show a significantly decreasing trend across the seven measured timepoints (**Fig. 6e**). These findings suggest that ex-vivo culturing of FSHD biopsies over time results in a phenotype that is more associated with “healthier muscle” - a result that is more striking in the more severely affected TA muscles compared to quadriceps.

Next, we tested the FSHD PhenoMapR signature in independent bulk^68–70^ and single-nucleus^71^ datasets. For the bulk datasets, the PhenoMapR signature derived from Wong et al.^61^ found significant stratification in two of three datasets (**Supplemental Fig. 7e**). In the single-nucleus dataset of primary TA muscle myoblast cultured cells, PhenoMapR showed significant separation between FSHD and control cells (**Fig. 6f**). Of note, PhenoMapR scores showed a significant positive correlation with the number of reported FSHD signature genes and FSHD signature level (**Fig. 6g; Supplemental Fig. 7f**). However, the elevated PhenoMapR score in many cells with low or undetectable FSHD signatures genes (**Fig. 6g**) suggests that PhenoMapR was able to detect disease-related populations that the reported signature would miss.

Magnetic resonance imaging (MRI) of FSHD-involved muscles has shown correlations between clinical presentation and MRI characteristics: samples with STIR MRI signal hyperintensity (STIR+) tend to have upregulated *DUX4* signatures compared to STIR- samples^70^. Using PhenoMapR to score bilateral TA muscle biopsies from patients with FSHD, we found a significant correlation between scores from paired bilateral biopsies (**Supplemental Fig. 7g**). Importantly, we also observed a significant enrichment for worse FSHD PhenoMapR scores in the STIR+ samples (**Fig. 6h**), as well as a significant correlation with STIR ratings at a more granular level (**Fig. 6i**). Interestingly, for samples with discordant STIR rating and PhenoMapR score (defined as low STIR rating ≤ 1 with high scaled PhenoMapR score > 0.5 (*n* = 5); circled points in **Fig. 6i**), high PhenoMapR score was associated with significantly higher inflammation (*p* ≈ 9×10⁻⁵), ECM (*p* ≈ 7×10⁻⁴), complement (*p* ≈ 0.02), *DUX4* target expression (*p* ≈ 0.016), and clinical severity score (CSS) (*p* ≈ 0.004) (**Supplemental Fig. 7h**). This shows that PhenoMapR successfully assigns samples along a known disease severity scale while also identifying disease-relevant signal not fully captured from standard clinical MRI measurements in FSHD.

Together, PhenoMapR was able to successfully identify disease-relevant samples and cells across FSHD bulk, single-cell, time course, and anatomical location data types from multiple independent datasets. These results broadly agree with known clinical measurement of disease severity (e.g. STIR rating) while also identifying low-risk patients that might be misclassified, as well as defining cell types and markers that might present therapeutic targets in FSHD.

## DISCUSSION

Previous phenotype mapping methods require a single bulk expression dataset as reference and lack the flexibility to incorporate precomputed signatures aggregated across multiple cohorts. PhenoMapR addresses both limitations: users can derive phenotype signatures from any labeled bulk dataset, or leverage PhenoMapR’s built-in PRECOG and TCGA meta-z scores (outcomes signatures aggregated across hundreds of bulk studies) that are robust to individual dataset variation by design.

We demonstrate PhenoMapR’s utility across bulk, single-cell, and spatial data in four biological domains. In COVID-19, PhenoMapR consistently identified neutrophil and monocyte populations as associated with severe disease and NK cells with milder disease across three independent large-scale single-cell cohorts. In cancer, PhenoMapR showed robust survival stratification across TCGA and, when applied to independent bulk, single-cell, and spatial PAAD datasets, identified stage-specific and stage-agnostic outcome associations, nominated canonical and unreported cell populations and drug targets in adversely prognostic cells, and revealed co-localization patterns and intercellular signaling axes between prognostic subpopulations. In Alzheimer’s disease, PhenoMapR identified phagocytically-impaired microglia as the most disease-associated cell type, a finding independently supported by large-scale AD meta-GWAS data. In FSHD, senescent vascular smooth muscle cells (VSMC) were most strongly associated with healthy phenotypes while metabolically stressed and proliferating VSMCs were tied to disease; notably, the FSHD phenotype signature validated in independent datasets, uncovered unique transcriptional features of ex-vivo cell culturing, and flagged disease-associated samples missed by established clinical measurements.

Several limitations warrant consideration. Like all phenotype mapping approaches, PhenoMapR’s performance depends on the quality and biological relevance of the reference signature used, an issue partially mitigated by the built-in multi-cohort meta-z functionality. The linear scoring model assumes additive gene contributions, which may not capture complex non-linear biology; however, we show this assumption produces negligible loss of biological signal relative to more complex methods, and its simplicity is precisely what enables scalability at atlas scale.

PhenoMapR provides a fast, scalable, and biologically accurate framework for mapping sample-level phenotypes onto single-cell and spatial transcriptomics data across any biological context where labeled bulk expression data exist. Compared to existing methods, PhenoMapR offers order-of-magnitude improvements in runtime and memory, pre-computed multi-cohort outcomes signatures, and, uniquely, an accompanying Shiny application that enables investigators without programming experience to perform the full analysis interactively. Together, these features position PhenoMapR as a broadly accessible tool for extracting phenotypic meaning from the increasingly large transcriptomic atlases now central to biological discovery.

## AUTHOR CONTRIBUTIONS

Brooks Benard: Conceptualization, Data curation, Formal analysis, Methodology, Project administration, Software, Supervision, Visualization, Writing—original draft. Chinmay Lalgudi: Software, Writing—review & editing. Armon Azizi: Conceptualization, Methodology. Andrew Gentles: Conceptualization, Funding acquisition, Project administration, Methodology, Supervision, Writing—review & editing.

## CODE AND DATA AVAILABILITY

The source code and test data for the PhenoMapR package and Shiny App are freely available online at Github (https://github.com/brooksbenard/PhenoMapR). Comprehensive documentation and vignettes are available at https://brooksbenard.github.io/PhenoMapR/index.html.

## DATA AVAILABILITY

All datasets included in this work are publicly available. Survival outcomes meta-z signatures for all cancers in TCGA and PRECOG are available at https://precog.stanford.edu/. Expression and outcomes data for TCGA were obtained from the OSF portal (https://osf.io/gqrz9/) and Genomic Data Commons (https://gdc.cancer.gov/about-data/publications/pancanatlas), respectively. COVID-19 single-cell datasets were obtained from public repositories: the Stephenson/Haniffa PBMC atlas as an AnnData file from CZI CELLxGENE (https://cellxgene.cziscience.com/); the Wilk et al. multi-omic COVID atlas as a Seurat RDS from the COVID-19 Cell Atlas (GSE174072; https://www.covid19cellatlas.org/index.html); and Schulte-Schrepping et al. as the public HCA AnnData corresponding to the Berlin 10x PBMC cohort (EGAS00001004571; Bonn Rhapsody data were not accessed). The preprocessed PAAD single-cell dataset CRA001160 and metadata were obtained from TISCH2^72^ (https://tisch.compbio.cn/home/). Single-cell and spatial data for the HTAN PAAD sample are available on the HTAN portal (https://humantumoratlas.org/). Bulk expression and phenotype annotations for AD datasets were obtained from the Gene Expression Omnibus using accession codes GSE39420, GSE109887, and GSE28146. Raw and processed data for the single-cell AD dataset are available at Zenodo (https://zenodo.org/records/17302976). The FSHA bulk and single-cell datasets were obtained from the Gene Expression Omnibus using accession codes GSE140261, GSE115650, GSE56787, GSE26852, GSE122873, GSE174301, GSE242912, and GSE143452.

## ACKNOWLEDGMENTS

We would like to thank all the authors of the included studies for making their data and code publicly available and useful to the scientific community. Special thanks to Dr. Ruo Han Wang, Dr. Amy Fan, Dr. Izumi de Los Rios Kobara, Dr. Brendan Ball, Louis Sharp, and Ilayda Ilerten for providing feedback on the PhenoMapR package and Shiny app.

## MATERIALS AND METHODS

### Gene expression preprocessing

PhenoMapR includes pre-computed meta-z gene signatures for outcomes from bulk transcriptomics datasets in TCGA and PRECOG^10^. These meta-z scores represent, for each gene in a disease (cancer) type, the strength of its statistical association with an outcome measurement - typically overall survival in the case of cancer. When a bulk expression dataset and phenotype labels are provided, datasets are processed in the same way as PRECOG, described as follows. For input expression data, reported gene IDs are updated to their currently approved HGNC HUGO IDs (package biomaRt v2.58.2). Any duplicate gene IDs are filtered by retaining the one with the greater average expression. Next, we (i) ensure the expression matrix is in log_2_ space, (ii) remove all genes with no variance (SD < 0.00001), (iii) remove genes/samples with ≥80% missing values, (iv) quantile normalize (for microarray datasets), (v) standardize (scale each gene to have mean zero and unit variance across samples), and (vi) impute missing values (for microarray datasets; package impute v1.76.0). Further details and motivation are provided in the original PRECOG study.

### Calculating bulk transcriptomics phenotype gene signatures

Univariate Cox proportional hazards regression is used to calculate gene-level z-scores for expression associations with time-to-event outcomes such as overall or progression-free survival in cancer (R package Survival v3.5–8). For inputs with binary phenotypes (e.g. responder/nonresponder; *n* < 5 per group throws warning), we used univariate logistic regression to calculate gene-level z-scores for expression associations with labels (R package Stats v4.3.1). For continuous phenotypes, we take the Pearson correlation between gene expression and numeric phenotype values, then convert to a z-score using a Fisher z-transformation (*`Fisher z-transform: z = 0.5 * log((1+r)/(1-r)) * sqrt(n-3))`*). Positive z-scores indicate that increased gene expression is associated with worse outcomes or the phenotype of interest (e.g. shorter survival time or time to progression, non-response to therapy, elevated cholesterol, etc.).

### Scoring of samples, cells, and locations

PhenoMapR maps phenotype signatures to samples, cells, and spatial locations using the weighted-sum between the phenotype signature (gene z-scores) and target expression matrix for the given input. The weighted-sum scoring approach allows more phenotypically-relevant genes to influence the aggregate score based on their sample/cell/location expression level. Additionally, this approach retains both absolute score and relative rank order of samples/cells/locations to allow for more dynamic analyses. For a given expression input (bulk, single-cell, or spatial), PhenoMapR first updates gene labels to currently-approved HGNC HUGO IDs and log-normalizes if needed. Then, using either the built-in TCGA/PRECOG meta-z signatures, user-provided phenotype gene z-score signature, or phenotype labeled bulk expression input z-score signature, we take the crossproduct of the expression matrix and the z-score gene signature (default filtered to genes with |Z ≥ 2|; R package crossprod v3.6.2).

Expression inputs may be dense or sparse matrices (genes × cells/spots), Seurat, SingleCellExperiment, SpatialExperiment, or AnnData objects. For Seurat/spatial objects, scoring uses the selected assay and layer (default normalized data; raw counts or scale.data are optional). Optionally, expression can be aggregated to pseudobulk profiles before scoring (pseudobulk = TRUE with a grouping column). Only genes present in both the query matrix and the filtered reference signature are retained. The PhenoMapR score for unit *j* is the weighted sum:

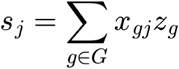

where *x_gj_* is expression of gene *g* in cell/spot *j*, *z_g_* is the reference association z-score, and *G* is the overlapping signature gene set (computed via a gene-aligned cross-product). For built-in overall-survival references, higher scores indicate more adverse phenotype association and lower scores more favorable association. An optional reference_sign multiplier (±1) can reverse score polarity without rewriting the reference matrix.

### COVID-19 single-cell ground-truth analysis

We established ground-truth benchmarking of PhenoMapR-identified cell-types using three large single-cell studies in COVID-19. For each cohort, cells (using author-provided labels) were split 50/50 within donor×cell-type strata. The training half formed donor pseudobulks (composition-insensitive all lineages included); the held-out half was scored with PhenoMapR references derived from those pseudobulks. Two phenotypes were fit: binary COVID vs healthy and continuous severity (ordinal; healthy = 0, asymptomatic = 1, mild = 2, moderate = 3, severe = 4, critical/fatal = 5). Cell-type importance was summarized by mean held-out scores and binary–continuous concordance. For each dataset, specific sample/condition filters were applied:

● Stephenson / Haniffa PBMC atlas: LPS / hospitalized non-COVID samples excluded.
● Schulte-Schrepping: only the Berlin 10x PBMC cohort was analysed since it was publicly-available; chemistry-matched healthy (10x 3′ v3 only) were used as controls to avoid v2/v3 confound (v2 controls were excluded).
● Wilk: analyses run separately on combined PBMC+WB and on PBMC-only and WB-only subsets to assess tissue effects on cell-type ranking.

### Activity-adjusted / cell-cycle–regressed scoring

The default mode (score_mode = "weighted_sum") applies the weighted sum to the input expression layer. Setting score_mode = "activity_adjusted" first regresses technical covariates from signature-gene expression using Seurat ScaleData, then computes the weighted sum on the residual (scaled) matrix. When S.Score and G2M.Score are among vars_to_regress, Seurat cell-cycle scores are computed with the updated 2019 S- and G2M-phase gene sets before regression. The default regressors are S-phase score, G2M-phase score, and library size (nCount_RNA); users may subset or extend this set (for example, cell-cycle scores alone). Library size is taken from metadata or computed as column sums when absent. This option is intended to reduce confounding of phenotype scores by proliferation and sequencing depth while preserving relative phenotype ranking.

### Empirical significance by gene-shuffle permutation

When permutation_n is set to a positive integer, PhenoMapR additionally reports an empirical *P* value for each cell/spot under a gene-shuffle null. Observed scores are computed with the true reference z-vector. For each of *N* permutations, reference z-scores are randomly reassigned among the same signature genes and scores are recomputed on unchanged expression, yielding a null distribution per unit. The empirical *P* for unit *j* is

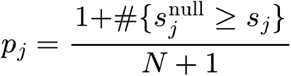

with a fixed random seed (permutation_seed, default 42). These *P* values assess whether a unit’s score is extreme relative to random shuffling of signature weights and expression.

### Donor-aware testing for CRA001160

To avoid treating cells as independent when they share a donor, we scored tumor cells from CRA001160 and aggregated to the donor. For each of 24 tumor donors we computed the median PRECOG score of each cell type, requiring ≥5 cells of that type in that donor. Ductal type 2 was then compared with every other cell type by a paired Wilcoxon signed-rank test (pairs = donors that had both types). *P* values were BH-adjusted across the 9 contrasts. The reported effect is the median of the paired differences (Ductal type 2 − other) with a bootstrap 95% CI. Stability was checked by leave-one-donor-out: dropping any single donor, Ductal type 2 remained the top-ranked type by mean of donor-medians in 24/24 iterations.

### Negative-control PRECOG scoring for CRA001160

Rank-order of cell type association with adverse outcomes in CRA001160 was tested using negative control signatures from unmatched cancers. The full CRA001160 tumor cohort was scored with PhenoMapR using PRECOG meta-Z references for Pancreatic (matched) versus Breast, Melanoma, Ovarian, Prostate, and Colon (mismatched), plus TCGA PAAD (*z*-cutoff = 2). For each reference, cell-type mean scores and ranks were compared, with emphasis on whether Ductal type 2 remained the top-scoring type under matched versus mismatched tissues.

### UMI downsampling

To test depth sensitivity, CRA001160 PAAD tumor cells were downsampled with Seurat SampleUMI from a 10,000 UMI/cell CP10k baseline to 75%, 50%, 25%, and 10% of baseline depth, then re–log-normalized and rescored with PhenoMapR (PRECOG Pancreatic; *z*-cutoff = 2). Stability was assessed by cell-level score concordance with full-depth scores and by preservation of cell-type median rank order.

### T-cell subset composition analysis in PAAD

To test whether adverse-associated T cells were specifically regulatory rather than a mixed exhausted or effector population, we restricted CRA001160 to tumor cells (41,976 cells) and scored them with PhenoMapR using the shipped PRECOG pancreatic reference. Phenotype extremes were the top and bottom 10% of scores across all tumor cells (no adverse T cells were detected with a 5% threshold), not a T-cell–only tail. T cells were then taken from the original Peng et al. annotation (celltype_original == "T"; *n* = 3,615), yielding 91 Most Phenotype+ (adverse) and 1,348 Most Phenotype− (favorable) T cells. A gene was called detected if its UMI count was > 0. Treg identity was defined as *IL2RA*⁺*FOXP3*⁺. Tregs were then split into activated Treg (Treg plus detection of *TIGIT*, *ICOS*, *CTLA4*, or *ENTPD1* / *CD39*) versus checkpoint-low Treg (Treg without those four genes). Remaining non-Treg T cells were assigned in mutually exclusive order: CD8⁺ (*CD8A* or *CD8B* detected, *CD4* not detected); CD4⁺ conventional (*CD4* detected, *CD8* not detected); CD4⁺CD8⁺; otherwise CD4⁻CD8⁻ / low lineage detect. Treg assignment overrode CD4/CD8 calls.

### Stratifying TCGA with PhenoMapR

Pre-processed gene expression data in TPM format for TCGA was downloaded from https://osf.io/gqrz9/^73^. For each TCGA cancer type with an equivalent PRECOG cancer, the expression matrix was first processed as previously described. Next, PhenoMapR was run on each cancer using the processed expression input and the matching PRECOG meta-z signature to derive sample-level scores. Using a median PhenoMapR score split within each TCGA dataset, we performed Cox-proportional hazards regression using TCGA outcomes data (Survival v3.5–8). Clinical outcomes data were obtained from https://gdc.cancer.gov/about-data/publications/pancanatlas^74^. Overall Survival data was used for all cancers except PRAD, DLBCL, LGG, READ, and TGCT, for which the Progression Free Interval (PFI) was used as recommended by Liu *et al.*^75^.

### Stage-specific signature generation and optimism-reduced evaluation in GSE253260

To assess whether stage-specific PhenoMapR signatures stratify overall survival (OS) beyond in-sample optimism, we performed a repeated train/test evaluation within the BACAP / GSE253260 PDAC cohort. Bulk RNA-seq expression and clinical annotations were obtained from GEO (GSE253260). Patients with available expression, clinical resectability stage (Resectable, Borderline, Locally Advanced, or Metastatic), and OS time/event status were retained. OS times recorded in days were converted to months when necessary. Analyses were conducted separately within each disease-stage stratum.

Within each stage, patients were randomly partitioned into a training set (70%) and a held-out test set (30%). On the training subset only, a gene-level Cox proportional hazards association signature was derived with PhenoMapR (`derive_reference_from_bulk()`), using OS time and event as the phenotype. Held-out patients were then scored with `PhenoMap()` using this stage-specific reference, retaining genes with absolute association |z| ≥ 2. Training labels were never used to score the test set. This procedure was repeated for 100 independent random splits per stage (fixed seeds for reproducibility). Stages with fewer than 20 patients, or splits yielding fewer than 10 training or 5 test patients, were excluded for that iteration.

On each held-out test set we evaluated prognostic discrimination and association as follows. The continuous PhenoMapR score was standardized within the test set (mean 0, SD 1), and a univariate Cox model of OS on the standardized score was fit to obtain a hazard ratio (HR) per 1 SD increase, Wald *p*-value, and Harrell’s concordance index (C-index).

### Cell type assignment for single cell and spatial datasets

We used cell-type annotations provided by the authors of the original studies when available. For the FSHD single-cell dataset, we used CyteTypeR^76^ to assign cell type labels since none were provided by the authors. Single cell data was downloaded from the Gene Expression Omnibus (GEO) using the accession ID GSE122873. Samples were combined and converted into a Seurat object using default settings (Seurat_5.4.0).

Using the top 10 positive cluster marker genes derived from FindAllMarkers() (log2 fold-change > 1), we ran CyteTypeR using default settings.

HTAN single cell and spatial data were downloaded from Synapse (https://www.synapse.org/). For the HTAN PAAD single-cell data (HT270P1-S1H2Fc2A2N1Bmn1_1scRNA_data.txt), cell types were identified using standard clustering, marker gene identification, and manual assignment of labels using known marker genes associations. Single-cell mapping of this sample to the paired ST sample (HT270P1-S1H2Fc2U1Z1Bs1-H2Bs2-Test) was performed using CytoSPACE (1.1.0; conda 23.1.0; solver method = lap_CSPR).

### Co-localization analysis of cell types in spatial transcriptomics data

Neighborhood co-occurrence analysis was performed on the CytoSPACE object for HT270P1-S1H2Fc2U1Z1Bs1-H2Bs2-Test using spatialCooccur (v 0.99.0) with 100 cell type label permutations for `nhood_enrichment()`. Cell types were labeled as either *CellType_N_*Adverse, *CellType_N_*Favorable, or *CellType_N_*Other based on their PhenoMapR score percentile (top and bottom 5th percentile were labeled as adverse and favorable respectively). The number of k-nn used was determined using `neighbors.k = min(20L, max(5L, nrow(scoc_df) - 1L)).`

### Cell-cell communication

Ligand–receptor signaling was inferred with CellChat v2^51^ on CytoSPACE-mapped cells from the HTAN PDAC Visium sample HT270P1-S1H2Fc2U1Z1Bs1-H2Bs2-Test. Cells were scored with PhenoMapR (PRECOG Pancreatic); phenotype extremes were the bottom 5% (Favorable) and top 5% (Adverse) of scores. CellChat was run on Adverse and Favorable cells only, labeled as {CellType}_{Adverse|Favorable}, after dropping labels with fewer than 10 cells (16 groups; 2,124 cells).

Expression was library-size normalized (CP10K, log1p). Communication probabilities used CellChatDB.human in spatial mode: truncated mean (trim = 0.1), distance-aware inference (distance.use = TRUE), secreted range 250 µm, contact/ECM range 100 µm, and filterCommunication(min.cells = 10). Interactions were aggregated by sender→receiver and stratified into physical (cell–cell contact + ECM–receptor) versus secreted signaling, then integrated with spatial neighborhood enrichment and local co-occurrence. Dual-supported pairs required both spatial co-localization (neighborhood *z* > 0 with BH *q* < 0.05, or local co-occurrence ≥ 0.5) and CellChat support (communication probability > 0 with ≥1 ligand–receptor pair).

### Marker genes for phenotype-associated cells

PhenoMapR provides the `define_phenotype_groups()` function that allows users to define the percentile of score extremes to use for defining “phenotype associated” cells (default = 5%). Once these groups are defined, PhenoMapR implements `find_phenotype_markers()` to identify cell-type agnostic or cell-type specific marker genes for cells/spots within the phenotype groups. For single-cell and ST data, `find_phenotype_markers()` implements Seurat’s `FindMarkers()` function (default settings). Markers can be selected by comparing against all other cells or against the same cell type in the opposite phenotype group. Results can be visualized as a heatmap using `plot_phenotype_markers().`

### Benchmarking PhenoMapR against other methods

We implemented SigBridgeR^35^, a toolkit that integrates 11 phenotype-mapping algorithms into a consistent workflow, to benchmark PhenoMapR in a rigorous approach. Using the same PDAC bulk (GSE253260) and single-cell datasets (CRA001160) used in Fig 3, we performed runtime and memory scaling analyses using subsets of 5k, 10k, 20k, and 50k cells. SigBridgeR default settings were used. We rank both survival and binary (primary and metastatic labels) implementations of each tool when available (SIDISH only provides survival phenotyping). To assess concordance with PhenoMapR across all methods, normalized enrichment scores were calculated for each phenotype label group using a gene set enrichment style approach against cells ranked by PhenoMapR score. Comprehensive cross-method comparisons were also performed. VmRSS polling was used to measure memory usage in order to capture both R and Python processes. Benchmarks were run on a Linux shared host (RHEL/Rocky 9–class kernel 5.14, x86_64) with two AMD EPYC 7413 24-core CPUs (48 physical cores / 96 threads, max ∼3.6 GHz) and ∼1.0 TiB RAM ( ∼4 GiB swap).

### DepMap CRISPR essentiality validation

To test whether PhenoMapR phenotype markers capture genes that are functionally important in pancreatic cancer cells, we compared CRA001160 malignant-ductal markers to CRISPR dependency profiles from DepMap. Single-cell phenotypes were scored on the full CRA001160 PAAD cohort (Peng et al.; TISCH2; 57,443 cells) using PhenoMapR with the PRECOG primary pancreatic signature. Phenotype extremes were defined as the top and bottom 10% of cells by PhenoMap score (in order to detect enough “favorable” ductal type 2 cells for analysis). Cell-type–specific adverse and favorable markers were identified with find_phenotype_markers using a within–cell-type contrast (`marker_scope` = "cell_type_specific", `celltype_contrast` = "within_cell_type", max_cells_per_ident = 5000). For DepMap comparisons we retained significant positive markers in malignant ductal epithelium (Ductal type 2) with FDR-adjusted *P* < 0.05 and average log₂ fold-change ≥ 0.5.

DepMap Public 26Q1 CRISPR screens were used (Chronos gene-effect and dependency-probability matrices). Because CRA001160 is a primary-tumor cohort scored with a primary PRECOG signature, models were restricted to primary PAAD lines (DepmapModelType == "PAAD" and PrimaryOrMetastasis == "Primary"; *n* = 25 lines with CRISPR data). Metastatic cultures were excluded.

For each gene, we computed the median Chronos gene effect and median dependency probability across retained models. Genes with median gene effect ≤ −0.5 were called essential; genes with median dependency probability ≥ 0.5 were called dependent. Marker–essentiality relationships were evaluated by (i) two-sided Fisher’s exact tests for overlap between marker sets and essential/dependent gene sets against the CRA001160 expression gene universe; (ii) Wilcoxon rank tests and one-sided Fisher tests for enrichment of markers in the most- or least-essential 10% tails of the genome-wide ranking; and (iii) gene-set enrichment analysis (fgsea; 10,000 permutations; gene-set size 10–5,000) on preranked essentiality ranks, treating Adverse and Favorable markers as pathways. Gene-effect and dependency modalities were analyzed separately for 2D PAAD lines and NextGen organoids.

### Orthogonal validation against scPANDA prognosis-related gene lists

PhenoMapR markers from CRA001160 were compared to published ctPANDA cell-type–resolved prognosis-related genes (Tang et al., *Cancer Cell* 2026; Table S2A), mapping worse-/better-survival genes to short-survival (SS) and long-survival (LS) sets. This is gene-list concordance only; the Tang expression atlas was not re-scored. Peng/TISCH labels were mapped to ctPANDA labels (e.g., Ductal type 2 → Ductal_PDAC). For each matched cell type we ran one-sided hypergeometric tests (CRA001160 genes as background; ≥5 genes per set) for Adverse–SS and Favorable–LS (expected) versus Adverse–LS and Favorable–SS (controls).

### Therapeutic target nomination using DGIdb

Adverse markers were identified as above (full CRA001160; PRECOG Pancreatic; 10% Most Adverse vs remaining cells within each cell type; FDR < 0.05; avg. log₂FC ≥ 0.5). Each gene×cell-type hit was annotated with: (i) DepMap primary-PAAD essentiality (median Chronos gene effect ≤ −0.5); (ii) membership in matched ctPANDA S2A SS sets and, where available, Table S4B worse-survival hazard ratios; (iii) housekeeping-like gene patterns; and (iv) a curated fallback druggable-gene list with illustrative drug hints.

Druggability was annotated using local DGIdb 5.0 flat files (Cannon et al., *Nucleic Acids Research* 2024; interactions.tsv and categories.tsv from https://dgidb.org/data/latest), not the legacy REST API. Per gene we summarized interaction counts, approved/antineoplastic/immunotherapy drug support, DGIdb categories (including CLINICALLY ACTIONABLE, DRUGGABLE GENOME, and cell-surface categories), and assigned tiers: A, clinical/oncology-supported DGIdb evidence; B, any drug–gene interaction; C, priority category without indexed interaction; D, curated list only; E, no support. Genes were considered DGIdb-tractable for novel-target panels if they were CLINICALLY ACTIONABLE or had ≥1 indexed interaction.

For cell-type–contextual nomination we further computed, within each focus cell type: specificity ratio (mean expression in the focus type / mean in all other types), expression mass fraction in the focus type, and Fav→Adv Δ*z* (difference in gene-wise *z*-scored expression between Most Adverse and Most Favorable cells of that type). Genes advancing to the novel/druggable panels were required to pass DGIdb tractability and expression gates of specificity ≥ 2×, Fav→Adv Δ*z* ≥ 0.75, and mass fraction ≥ 10%, and were ranked by trajectory score Δ*z* × log(1 + specificity). A composite priority score additionally weighted DepMap essentiality, cell-type uniqueness, ctPANDA SS support, elevated S4B hazard ratio (HR ≥ 1.5), DGIdb tier, non-housekeeping status, malignant-ductal context, and cell-surface annotation. Shortlists emphasized DepMap-essential, non-housekeeping, DGIdb- or curated-druggable adverse markers, with separate views for cell-type–unique and ctPANDA-supported candidates.

### Alzheimer’s disease meta-GWAS validation

We derived a custom bulk reference from GSE39420 (Early AD vs control; *n* = 7 vs 7) with derive_reference_from_bulk, scored a brain snRNA-seq cohort (GSE138852, https://zenodo.org/records/17302976) with PhenoMapR (`z_score_cutoff` = 2), and defined phenotype extremes as the top and bottom 5% of scores. Cell-type–stratified markers were identified with find_phenotype_markers (Wilcoxon; FDR-adjusted *P* < 0.05; avg. log₂FC ≥ 0.25), contrasting Most Adverse vs Most Favorable within each cell type (Mode 2: vs opposite tail). Significant markers were tested for enrichment of Alzheimer’s risk genes from the EADB–IGAP–PGC consensus meta-analysis (Bellenguez et al., *Nat. Genet.* 58:1214–1225, 2026; Supplementary Table 2), using tier 1 and tier 2 loci (109 loci; 139 unique gene symbols after locus expansion). Enrichment used one-sided Fisher’s exact tests against the full expressed gene background, with Benjamini–Hochberg correction within each tier subset.

### Validation of PhenoMapR Alzheimer’s signature in independent datasets

The Early AD–derived PhenoMapR signature (ad_reference from GSE39420; Early AD vs control via `derive_reference_from_bulk`) was applied to two independent GEO bulk cohorts downloaded with GEOquery. Probe-level expression was collapsed to gene symbols (mean of probes mapping to the same symbol), and each sample was scored with PhenoMapR using the same reference (`z_score_cutoff` = 2). GSE109887 samples were labeled Control vs AD from GEO metadata. Score differences between groups were tested with a two-sided Wilcoxon rank-sum test. GSE28146 samples were assigned to ordered stages (Control, Incipient, Moderate, Severe) from metadata text. Overall stage effects were tested with a Kruskal–Wallis test; pairwise Control vs stage contrasts used Wilcoxon tests for visualization. For plotting, scores were *z*-scaled within each cohort.

### Donor variation in Alzheimer’s microglia ranking

AD single-cell profiles were scored with a bulk-derived AD reference from GSE39420, then aggregated to donor×cell-type median scores (donors with ≥5 cells per type). Microglia were compared to other cell types with paired Wilcoxon tests across donors (bootstrap CIs on paired median differences), leave-one-donor-out stability of the top rank, and within-donor cell-type label permutation to assess whether the microglia advantage depended on donor structure.

### FSHD external validation

All three analyses scored samples with the same custom PhenoMapR signature (ref_fshd) derived from GSE140261 bulk muscle biopsies (27 FSHD + 8 controls; *Wong et al.*, *Hum. Mol. Genet.* 2020) using `derive_reference_from_bulk` (binary FSHD vs. control; higher score = more FSHD-like). Scoring used PhenoMapR with `z_score_cutoff` = 2.

TA vs quadriceps (GSE174301). Primary FSHD myoblast RNA-seq (RSEM TPM; *n* = 36: 8 TA, 28 Quad) was scored and globally *z*-scaled. Samples were stratified by muscle (tibialis anterior vs quadriceps) and day of differentiation. TA vs Quad scores were compared within Day 0 and Day 12 by Wilcoxon tests (one-sided TA > Quad); Day 0 vs Day 12 changes within each muscle used Wilcoxon tests; Quad score decline across differentiation days was tested with a Jonckheere–Terpstra test (alternative = "decreasing").

scRNA Control vs FSHD (GSE122873). Single-cell RNA-seq from Van den Heuvel *et al.* (*Hum. Mol. Genet.* 2019; 4 FSHD + 2 control cultures; 6,898 QC-passing cells after removing cells with ≥10% mitochondrial reads) was processed in Seurat (normalization, PCA, clustering, UMAP), scored with ref_fshd, and compared Control vs FSHD globally and within cell types by Wilcoxon tests. Phenotype extremes (top/bottom 5% of scores) were used for marker discovery with `find_phenotype_markers` (FDR < 0.05).

STIR MRI association (GSE242912). Bilateral tibialis anterior biopsies (*n* = 64; Wellstone clinical annotations) were CPM-normalized, scored with ref_fshd, and globally *z*-scaled. STIR+ vs STIR− scores were compared by Wilcoxon test. Association with ordinal STIR rating (0–4) used Spearman correlation. Left–right score concordance across paired biopsies (*n* = 31 subjects) was assessed by Pearson correlation. We flagged STIR–score discordant biopsies as those with STIR rating ≤ 1 but globally scaled PhenoMapR score > 0.5 (*n* = 5), i.e. transcriptionally FSHD-like samples with little or no STIR MRI signal. The reference group was remaining low-STIR biopsies with scaled score ≤ 0.5 (*n* = 41). Discordant vs reference biopsies were compared on Wellstone clinical and transcriptional summary features (inflammation, ECM, complement, DUX4-target programs, pathology cumulative score, MRI fat fraction, CSS) using two-sided Wilcoxon rank-sum tests.

### PhenoMapR Shiny Application

PhenoMapR includes an interactive Shiny application that exposes the package workflow without requiring users to write R code. The app is shipped with the package (`inst/shiny`) and launched locally with `PhenoMapR::run_app()`. It is implemented in R (v4.5.2) using shiny (v1.14.0) and bslib (v0.11.0, Bootstrap 5), with interactive tables and plots rendered via DT (v0.34.0) and ggplot2 (v4.0.3).

The interface is organized as a sequential, tabbed workflow that mirrors the package API: (1) upload expression data and optional cell metadata; (2) select a built-in phenotype reference (PRECOG, TCGA, Pediatric PRECOG, or ICI-PRECOG) or supply/derive a custom signature; (3) compute continuous phenotype scores; (4) visualize score distributions and embeddings or spatial layouts; and (5) define phenotype extremes and identify marker genes. Supported inputs include Seurat, SingleCellExperiment / SpatialExperiment, dense or sparse matrices, 10x HDF5, and AnnData (.h5ad).

Computationally, the app is a thin UI layer over the same exported functions used programmatically (`PhenoMap()`, `define_phenotype_groups()`, `find_phenotype_markers()`, `derive_reference_from_bulk()`, and related diagnostics), so interactive and script-based analyses are equivalent. All computation runs on the user’s machine (or a local/HPC host); no data are sent to an external server.

## SUPPLEMENTARY FIGURES

**Supplemental Fig. S1:**
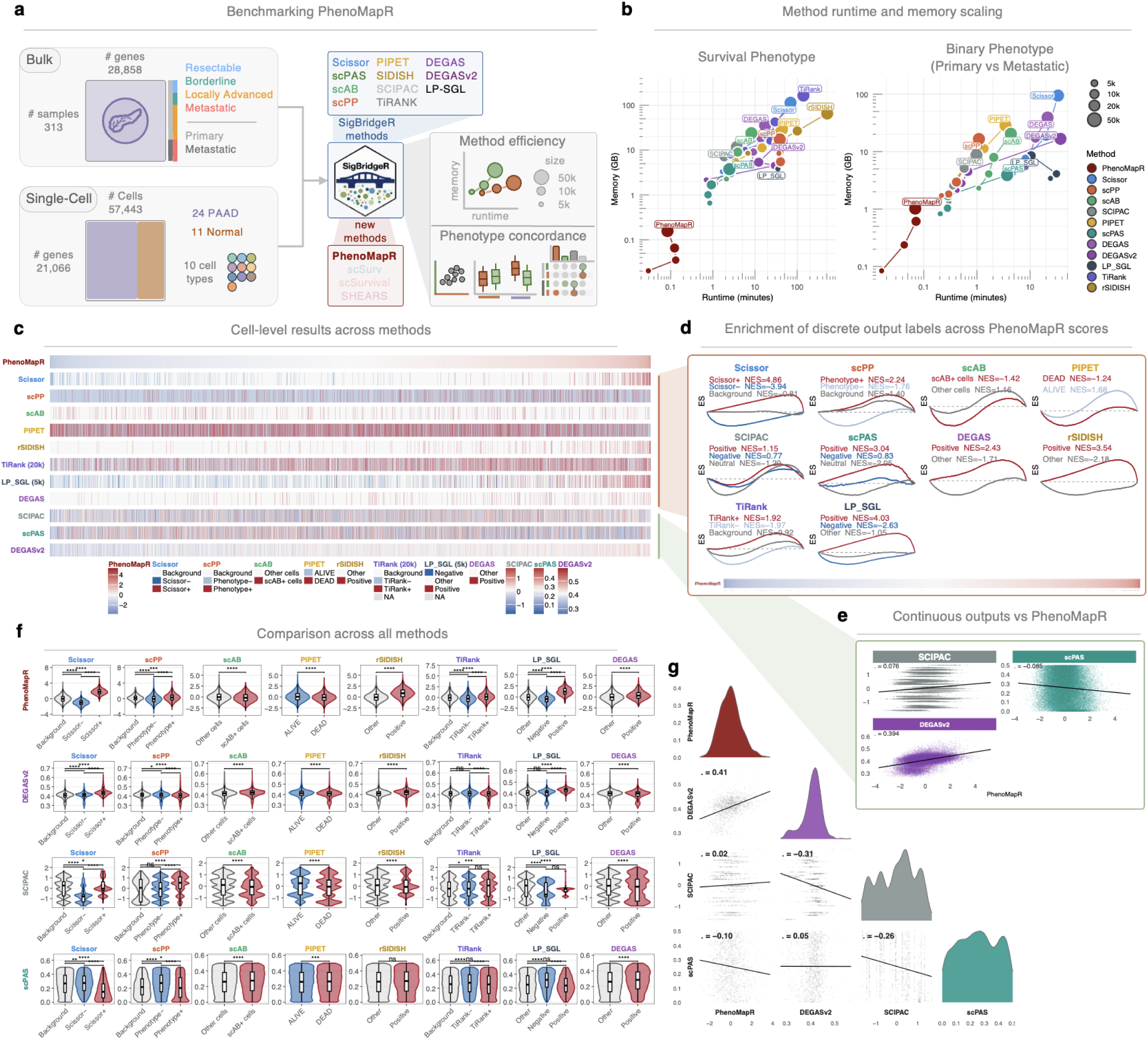
Benchmarking of PhenoMapR. **(a)** Benchmarking approach using SigBridgeR across twelve methods to identify phenotype-associated cells from bulk and single-cell PAAD datasets. **(b)** Memory and runtime scaling across 12 methods that map sample phenotypes to single-cell data. All measurements are taken around the core function call of each method after all preprocessing has been performed by SigBridgeR. **(c)** Cell-level results for all methods tested, ranked by PhenoMapR score. **(d)** Enrichment score plots for methods with discrete cell label results. Each plot is ranked by PhenoMapR score and the enrichment of phenotype-associated cells for each method are measured relative to the PhenoMapR rank. **(e)** Scatterplots between PhenoMapR results and results for methods that generate continuous outputs. **(f)** Cross-method comparison of scores derived from methods with continuous results (y-axis) to methods with discrete results (x-axis). **(g)** Correlation analyses between all methods that generate continuous scores.

**Supplemental Fig. S2:**
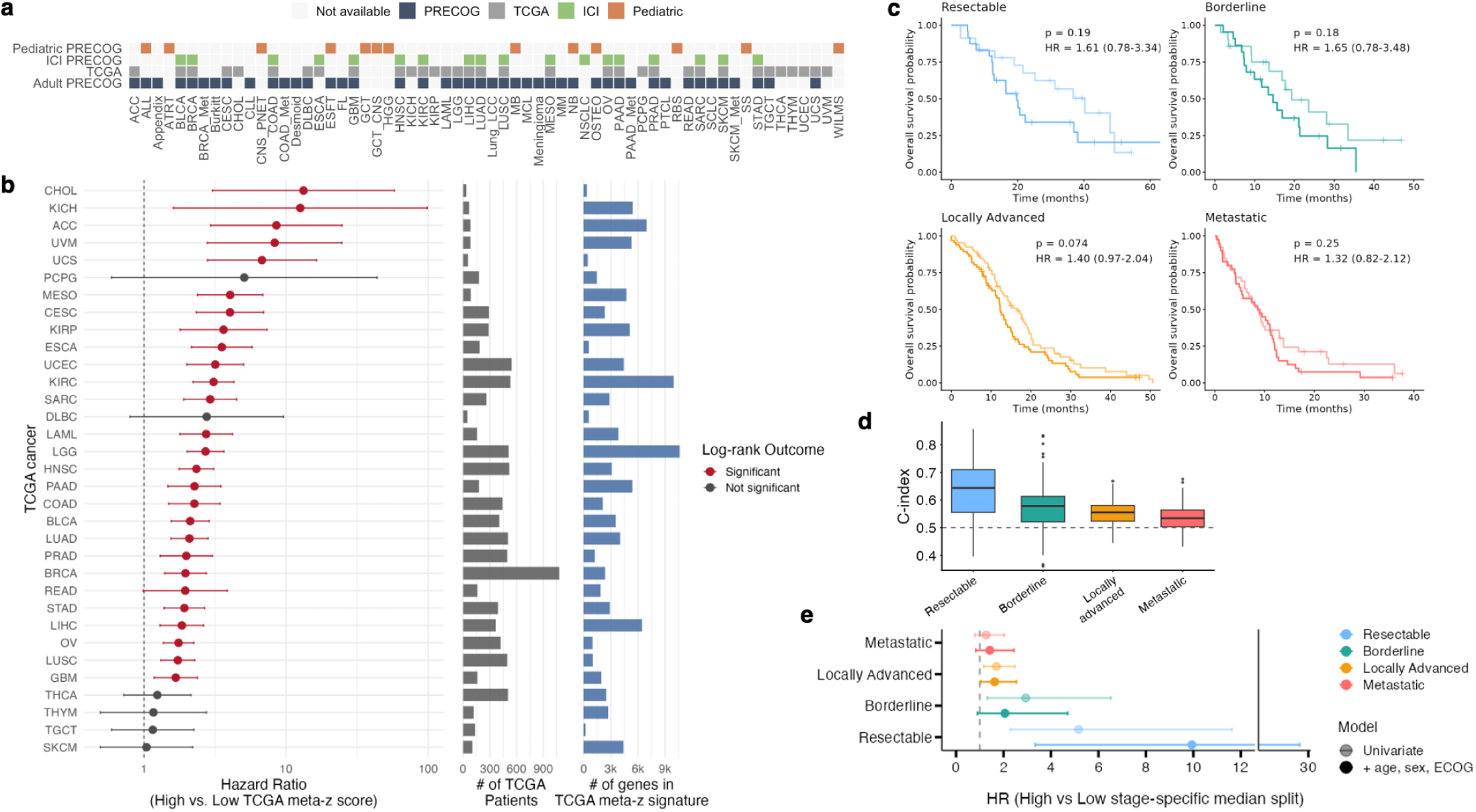
Validating PhenoMapR on bulk gene expression datasets. **(a)** Built-in PhenoMapR gene meta-z scores across TCGA and PRECOG. PhenoMapR contains pre-computed meta-z signatures for outcomes associations across adult, pediatric, and immunotherapy cohorts in PRECOG and TCGA. **(b)** Forest plot of Hazard Ratios across TCGA cancer types calculated using a median PhenoMapR score split. Scores were generated using TCGA meta-z scores. **(c)** Distribution of age, sex, ECOG status, and disease stage across the GSE253260 PAAD cohort. **(d)** Kaplan-Meier curves for the four disease stages split by median full-cohort PhenoMapR score within each stage. **(e)** Held-out concordance of stage-specific PhenoMapR signatures in GSE253260. Boxes show the distribution of Harrell’s concordance index (C-index) from a univariate Cox model of the continuous signature score on the test set; the dashed line marks C = 0.5 (no discrimination). Higher C-index indicates stronger ranking of overall survival risk on unseen patients within that stage. **(e)** Forest plot of univariate and multivariate hazard ratios within each stage-specific signature scoring.

**Supplemental Fig S3:**
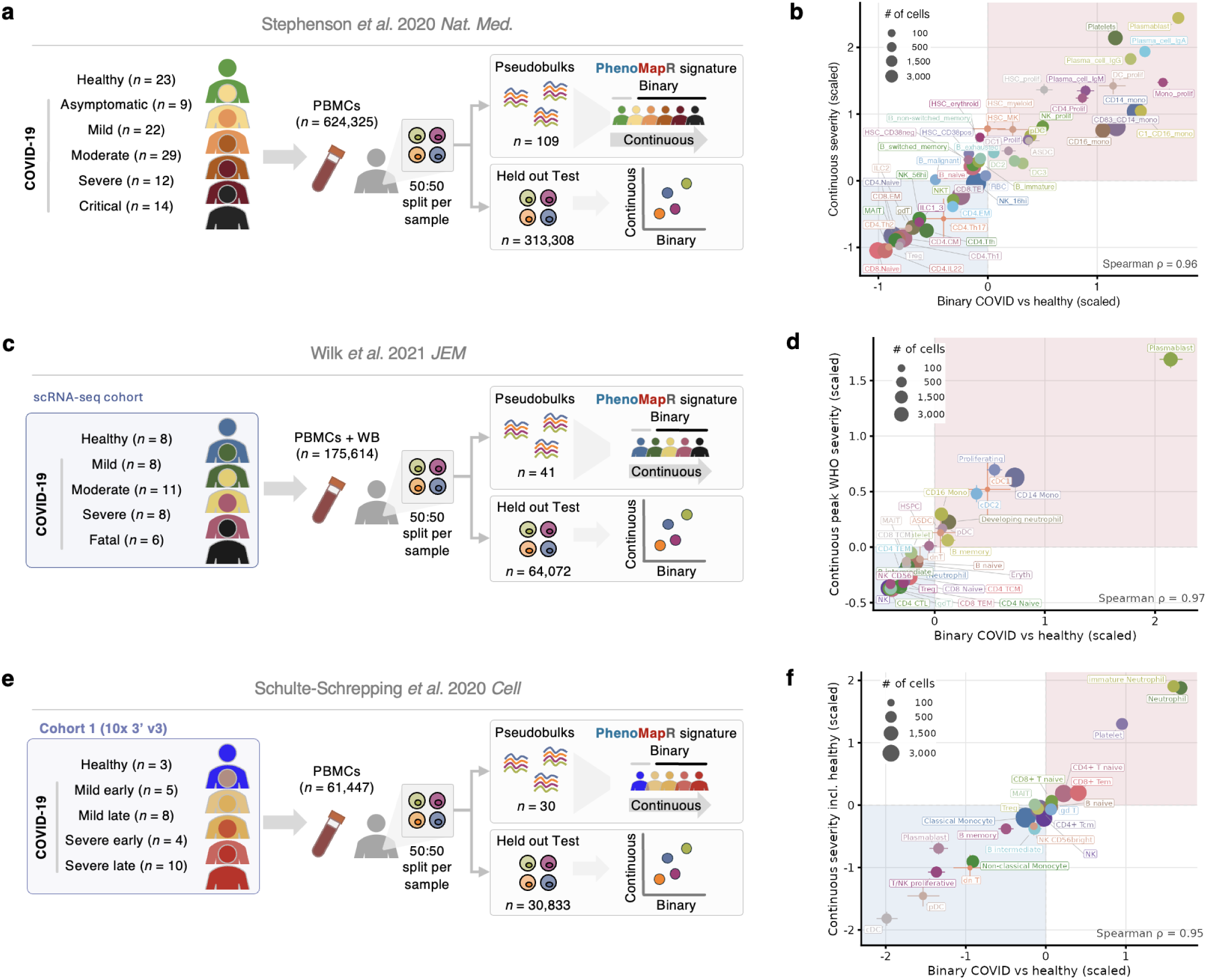
Positive control analyses using pseudobulk signatures from single-cell COVID-19 datasets. **(a)** Schematic depicting the disease severity spectrum and sample pseudobulk (train) vs. single-cell (test) split in COVID-19 patients from the Stephenson dataset. **(b)** Scatterplot of continuous (y-axis) vs. binary (x-axis) scaled PhenoMapR scores for each cell type in the Stephenson dataset. **(c)** Schematic depicting the disease severity spectrum and sample pseudobulk (train) vs. single-cell (test) split in COVID-19 patients from the Wilk dataset. **(d)** Scatterplot of continuous (y-axis) vs. binary (x-axis) scaled PhenoMapR scores for each cell type in the Wilk dataset. **(e)** Schematic depicting the disease severity spectrum and sample pseudobulk (train) vs. single-cell (test) split in COVID-19 patients from the Schulte-Schrepping dataset. **(f)** Scatterplot of continuous (y-axis) vs. binary (x-axis) scaled PhenoMapR scores for each cell type in the Schulte-Schrepping dataset. Error bars for all scatterplots represent the 95% bootstrapped confidence interval of each cell type’s mean score. Red and blue regions reflect shared quadrants of more severe (red) and less severe (blue) clinical disease.

**Supplemental Fig. S4:**
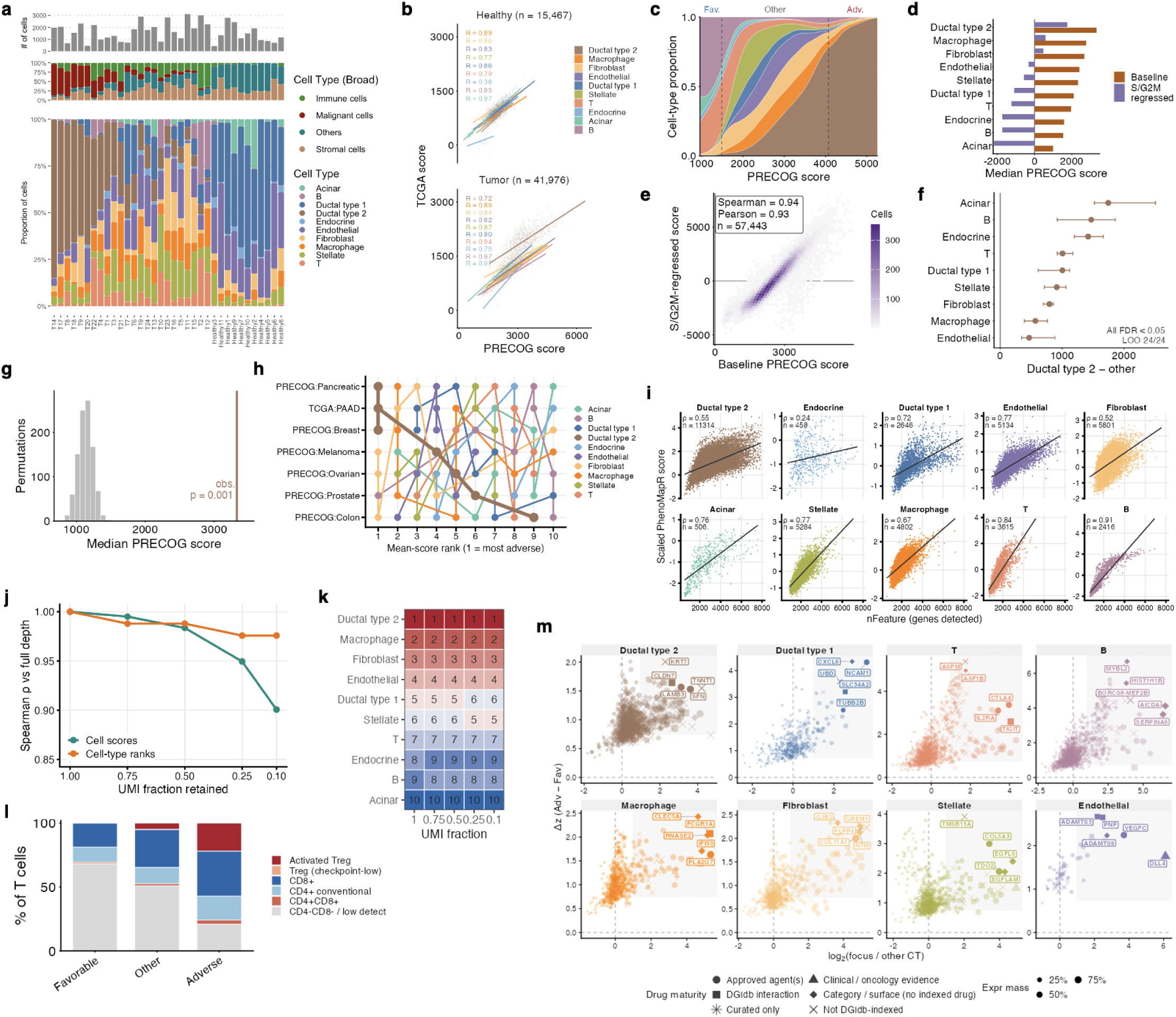
Applying PhenoMapR in a single-cell PAAD dataset. **(a)** Number and proportion of cells per cell-type and sample (left) along with their UMAP overlay (right). **(b)** Scatterplots between all single-cell PhenoMapR scores using TCGA or PRECOG meta-z references across healthy and tumor samples. **(c)** Proportion of cell types (tumor only) across the range of PRECOG PhenoMapR scores. Dashed lines indicate the thresholds for the upper and lower 5^th^ percentile of scores. **(d)** Bar plot for median score per cell type with or without regression of cell-cycle programs (S/G2M), ranked by median non-regressed cell type scores. **(e)** Scatterplot and correlations between baseline and cell-cycle regressed PhenoMapR scores across all cells. **(f)** Observed Ductal type 2 median score (brown line) versus the shuffled PRECOG signature weights distribution. The observed value lies well above the null (*p* = 0.001). **(g)** Within-donor difference in median score between Ductal type 2 and each other cell type (points, 95% CI). All contrasts remain significant after FDR correction, and leave-one-out donor cross validation. **(h)** Rank-order of mean cell type prognostic scores using different non-matched cancer meta-z signatures from PRECOG. **(i)** Scatterplots of scaled PhenoMapR scores and the number of genes detected per cell. Correlation calculated using Spearman rho. **(j)** Effect of down sampling UMIs on cell scores and cell-type ranks compared to full depth scores. **(k)** Cell-type rank-order based on median PhenoMapR score at different down sampling fractions. **(l)** Stacked barplot for lineage composition of all tumor T cells split by 5^th^ percentile thresholds. **(m)** Scatterplots of top adverse marker genes across cell types based on expression delta between within cell type adverse vs. favorable expression (y-axis) and cell-type specific expression versus all other cell types (x-axis). Points are shaped based on clinical actionability as categorized by DGIdb. Points are sized based on “expression mass” which is defined as the summed within cell-type expression divided by the summed expression across all other cell types.

**Supplemental Fig. S5:**
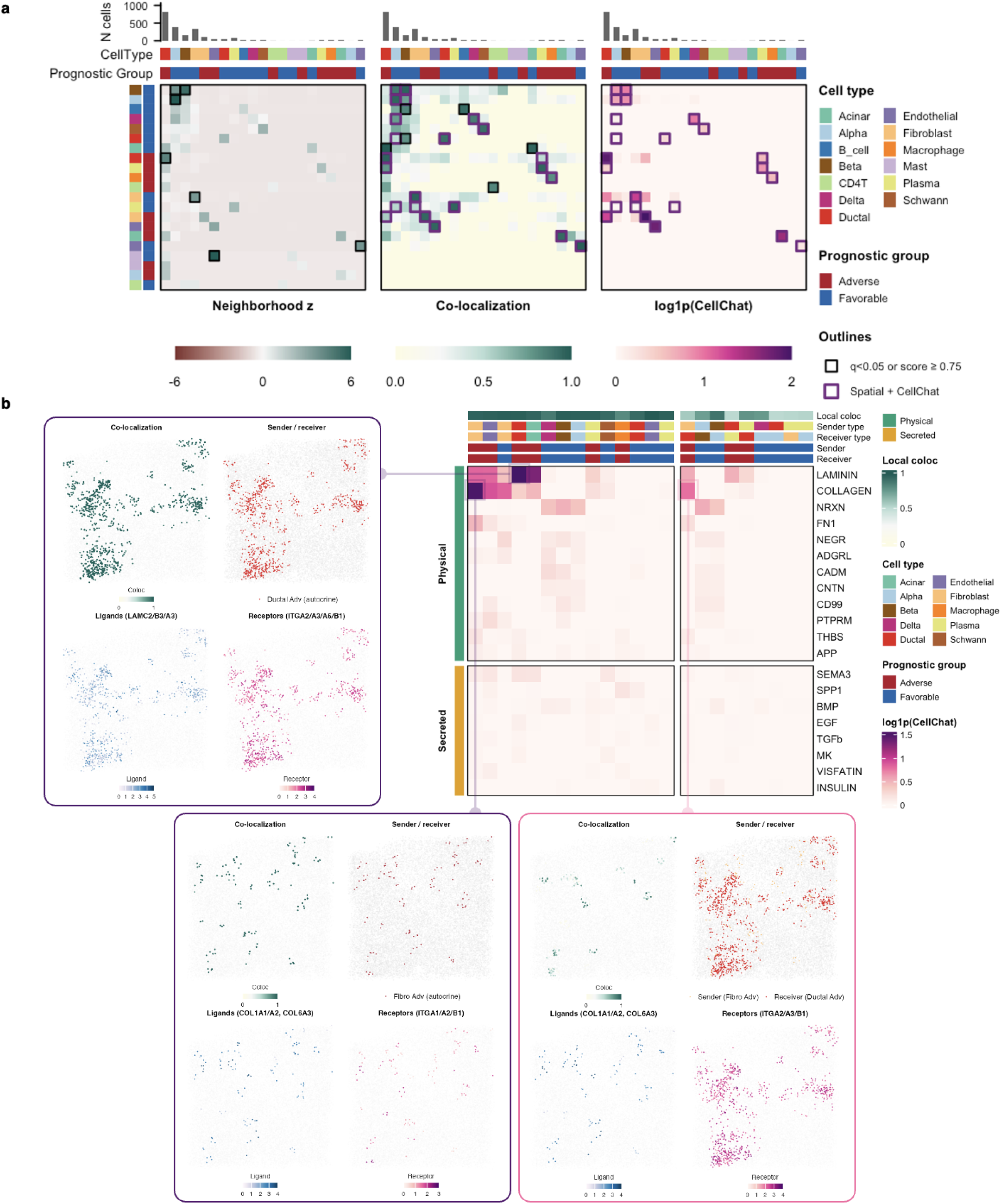
Spatial correlates of prognostic cell co-localization and cell-cell communication. **(a)** Heatmaps of neighborhood enrichment (left), local co-localization (middle), and cell-cell communication potential (right). **(b)** Summary heatmap and examples of top communication axes across prognostic cell populations.

**Supplemental Fig. S6:**
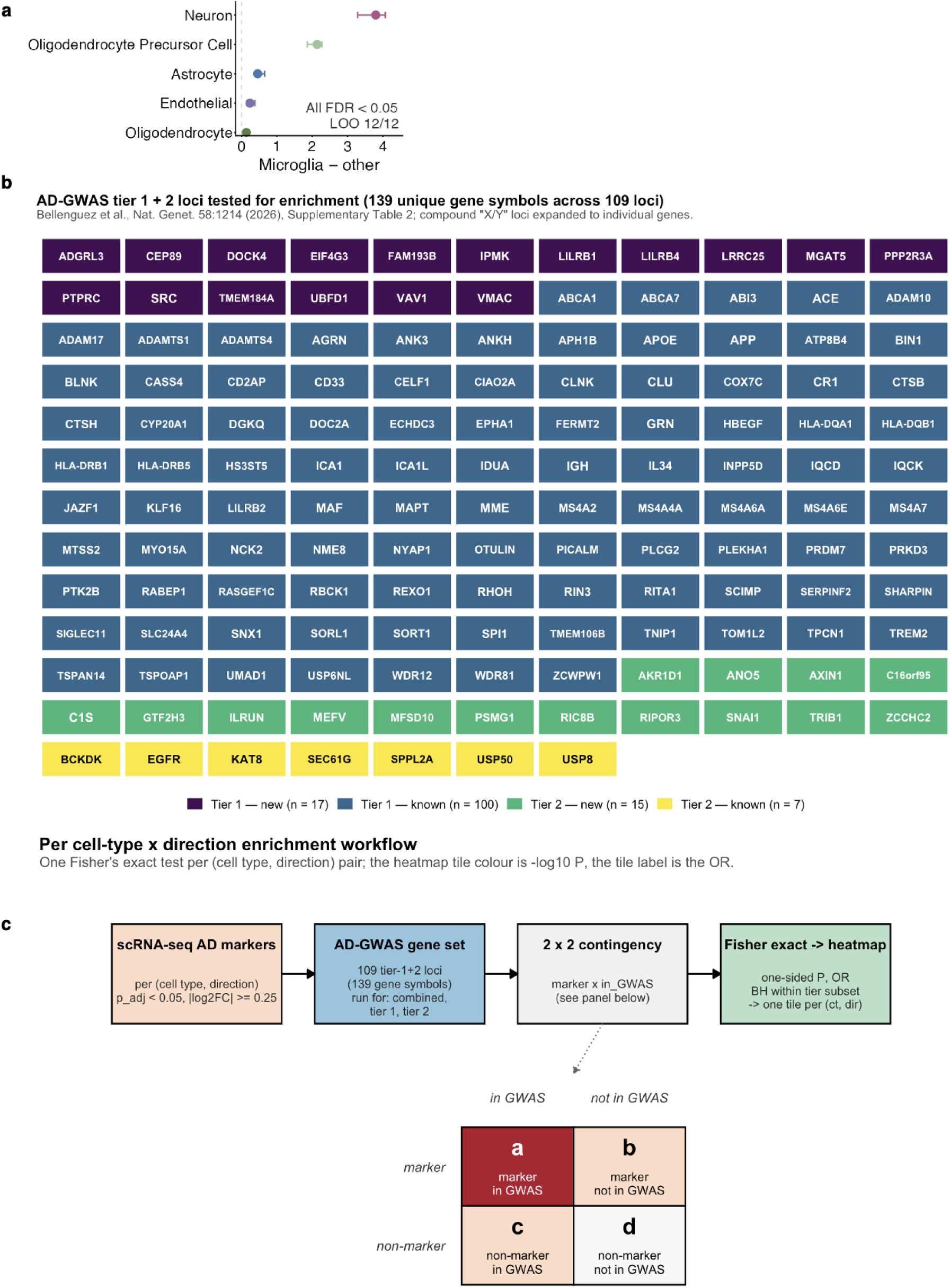
Meta GWAS genes tested for enrichment in Alzheimer’s disease. **(a)** Within-donor difference in mean score between microglia and each other cell type (points, 95% CI). All contrasts remain significant after FDR correction, and leave-one-out donor cross validation. **(b)** Table of genes reported as strongly linked to Alzheimer’s disease development. Color is based on reported “tiers” and previously reported or novel status. **(c)** Workflow of testing for enrichment of meta GWAS genes in the cell-type marker genes identified by PhenoMapR to be associated with Alzheimer’s disease.

**Supplemental Fig. S7:**
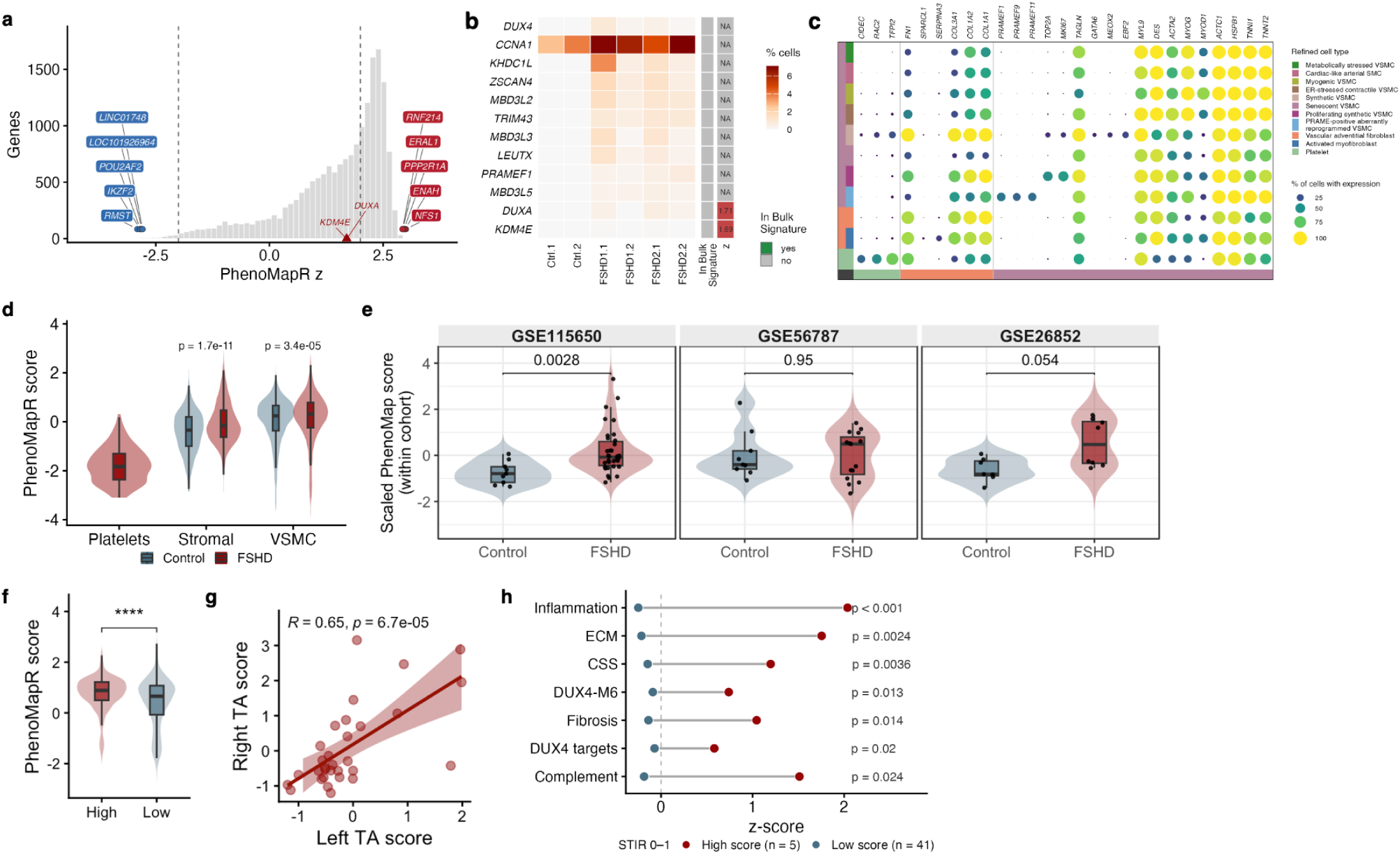
Single-cell and bulk expression associations between PhenoMapR scores and FSHD. **(a)** Distribution of bulk PhenoMapR derived FSHD signature. Top and bottom five genes by z-score are labeled. Genes labeled with red triangles reflect canonical *DUX4* target genes. **(b)** Heatmap of canonical *DUX4* target gene expression across single cell samples and whether they are present in the bulk FSHD signature. **(c)** Marker gene bubble plot for CyteTypeR defined cell types in the FSHD single-cell dataset. **(d)** PhenoMapR score distribution based on broad cell types and sample type. **(e)** Distribution of PhenoMapR scores between control and FSHD samples across three bulk expression datasets. **(f)** Distribution of PhenoMapR scores by the number of FSHD signature genes detected per single cell. **(g)** Scatterplot of PhenoMapR scores for bilateral TA muscle biopsies from FSHD patients. **(h)** Enrichment of biological signatures and clinical measurements in samples with discordant PhenoMapR scores and STIR ratings.

